# Biophysical Modeling Elucidates Mechanistic Principles for Rational Molecular Glue Design

**DOI:** 10.64898/2025.12.24.694639

**Authors:** Seokjoo Chae, Jonathon DeBonis, Joseph Quinlan, Ryan Richards, Junmin Wang, Gustavo Gutierrez, Jonathan Tart, Maxime Couturier, Sergio Martinez Cuesta, Meizhong Jin, Oleg A. Igoshin

## Abstract

Molecular glues are small molecules that offer a powerful strategy to target previously “undruggable” proteins of interest (POI) by enhancing their interactions with other proteins (effector). Depending on the nature of the effectors, molecular glues can induce stabilizing sequestration or degradation of POI. However, their rational design has been hindered by a poor understanding of how kinetic parameters impact their performance. To address this, we developed a unified mathematical framework that accurately analyzes the dynamics of both glue degraders and stabilizers. Our model reveals that the strong binding affinity of the ternary complex is the key determinant of performance across both modalities, provided that the initial component concentrations and degradation rate constants are fixed. Furthermore, we demonstrate that degrader performance is ultimately limited by its catalytic efficiency and the target protein’s natural half-life. We also identify distinct roles for effector abundance, showing that the relative concentrations of the effector and POI are critical for stabilizers but less so for degraders. This quantitative framework provides mechanistic principles for the rationale design and optimization of molecular glues.

## Introduction

Chemical-induced proximity (CIP) is a powerful approach for disease treatment that creates non-native protein-protein interactions (PPIs), enabling a range of downstream effects. CIP-based drugs, such as proteolysis-targeting chimeras (PROTACs) and molecular glues (MGs), elicit therapeutic effects by inducing non-native PPIs between an effector protein and a target protein of interest (POI). These drugs most commonly induce the inhibition or degradation of pathogenic proteins; however, recent advances have also shown promise in applications such as transcriptional activation as well.^1–3^ While PROTACs -- heterobifunctional molecules with specific binding moieties connected by a linker -- have dominated the CIP landscape due to their engineerable, modular design, they often suffer from poor drug-like properties, including limited oral bioavailability and cellular uptake, and exhibit a hook effect that further complicates exposure-response relationships.^4–6^ MGs, as monovalent compounds, offer more attractive drug-like properties.^7^ However, unlike the rational design approaches available for PROTACs, MG discovery has historically depended on serendipitous observations. While high-throughput screening approaches have recently shown promise in identifying MGs, the mechanistic principles governing how specific kinetic parameters translate to therapeutic outcomes remain elusive^4,5^. Thus, establishing a predictive framework to guide rational MG design is an urgent and unmet need in the field.

Depending on the nature of the effector proteins, MGs can induce the stabilizing sequestration (MG stabilizers) or degradation (MG degraders) of POIs. MG stabilizer examples include fusicoccin and related compounds that stabilize the interaction between 14-3-3 proteins and the POI, which have been used as tools to study and modulate such interactions^8,9^. MG degraders, in contrast, induce the degradation of the POI through mediating ternary complex formation with E3 ubiquitin ligases, subsequently tagging the POI for proteasomal destruction through ubiquitination. The most clinically successful MG degraders examples are the immunomodulatory drugs (IMiDs) such as thalidomide, lenalidomide, and pomalidomide, which bind to the E3 ligase cereblon (CRBN) and recruit POIs such as IKZF1 and IKZF3 for degradation, making them effective treatments for multiple myeloma and other hematological malignancies^2,3^. To understand how kinetic parameters affect these diverse MG dynamics, we need a universal framework that comprehensively covers all MG mechanisms.

Existing mathematical models of CIP in the literature have either focused on the PROTAC modality^10,11^ or on developing a pipeline for MG characterization^12^. While nicely providing a mechanistic framework for representing small-molecule-induced ternary complex formation with or without subsequent degradation, a comprehensive analysis of how specific model parameters relate to MG performance is lacking. In this work, we provide detailed insight into the relationship between MG kinetic parameters and performance by developing and validating mathematical models of four distinct MG systems and analyzing the effects of perturbing model parameters on predicted MG performance. Our results are the first to directly investigate these relationships and provide a quantitative rationale for the engineering of MG parameters.

## Methods

### Mathematical Model of FKBP Stabilizer

The dynamics of the FKBP stabilizer are modeled based on a two-step binding process. First, the MG stabilizer (G) binds to the effector protein, FKBP12 (E). Subsequently, this binary complex (EG) binds to the FRB domain of the POI, mTOR (P), to form a ternary complex (EGP).

This system can be described at equilibrium by the following algebraic equations, based on the law of mass action:

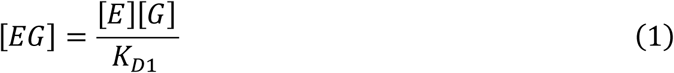

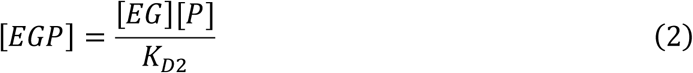

Where:

- *[E], [G]*, and *[P]* are the concentrations of the free effector (FKBP12), MG stabilizer, and target protein (mTOR), respectively (nM).
- *[EG]* is the concentration of the binary complex (nM).
- *[EGP]* is the concentration of the ternary complex (nM).
- *K_D1_* is the dissociation constant for the formation of the binary complex (nM).
- *K_D2_* is the dissociation constant for the formation of the ternary complex from the binary complex (nM).

The total concentrations of each species are constrained by the following mass balance equations:

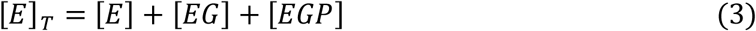

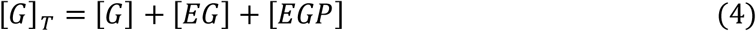

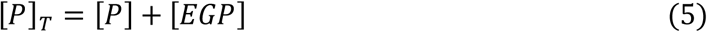

By solving this system of equations, the steady-state concentrations of each species can be determined. From these values, the normalized signal for a fluorescence polarization assay can be calculated as the fraction of the total effector protein (*E*_*T*_) that has formed the ternary complex:

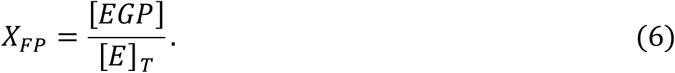

### Mathematical Model of CYPA Stabilizer

To describe the dynamics of the CYPA stabilizer (G), its target protein (P), and the enzyme (E, presumably CYPA), we utilized the following system of ordinary differential equations:

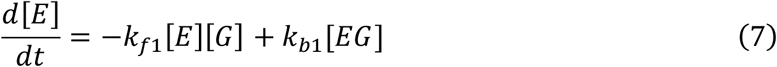

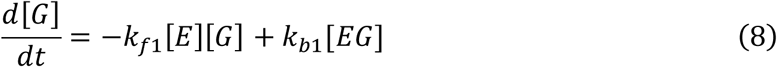

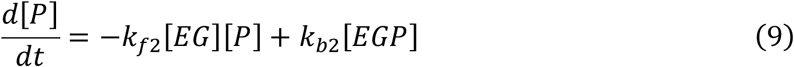

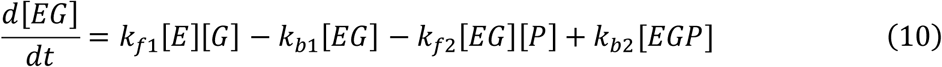

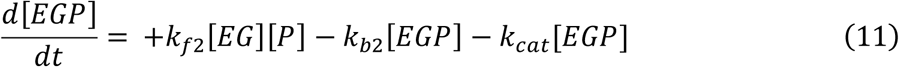

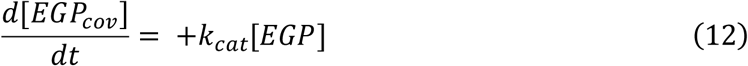

Where:

- *[E], [G]*, and *[P]* are the concentrations of the free enzyme, MG, and target protein, respectively (nM).
- *[EG]* is the concentration of the binary effector protein:MG complex (nM).
- *[EGP]* is the concentration of the non-covalent ternary effector protein:MG:POI complex (nM).
- *[EGP_cov_]* is the concentration of the final covalent complex (nM).
- *k_f1_* and *k_b1_* are the forward and backward rate constants for the *E* + *G* ↔ *EG* binding, and they are assumed to be *10^5^M^-1^s^-1^*.
- *k_f2_* and *k_b2_* are the forward and backward rate constants for the *EG* + *P* ↔ *EGP* binding (nM/s).
- *k_cat_* is the rate constant for the irreversible covalent modification step *EGP* → *EGP*_*cov*_ (1/s).

The TR-FRET signal (*X_TRF_*) was assumed to be proportional to the total concentration of the ternary complex (both non-covalent and covalent). The simulated signal was normalized to a 0-100 scale, where the signal at the maximum MG concentration (*G_max_*) was set to 100.

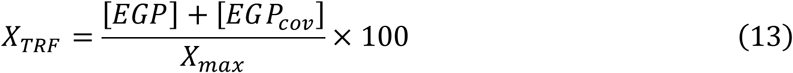

Where *X_max_=[EGP]+[EGP_cov_]|_G=Gmax_*.

The covalent modification assay was simulated by tracking the depletion of the non-covalent target protein species over time. The fraction *f*_*nc*_ of total protein concentration (*P_T_*) that is not in a covalently-bound complex (i.e., remaining as either as free protein, *[P]*, or as part of the non-covalent complex, *[EGP]*) was calculated.

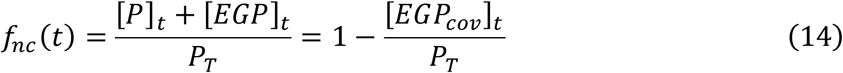

This decay curve was fitted to a single exponential function, *exp(-X_cov_t)*, to extract the observed decay rate constant, *X_cov_*.

### Mathematical Model of 14-3-3 Stabilizer

The dynamics of the 14-3-3 stabilizer is modeled based on a system where the ternary complex can form through two distinct pathways. The effector protein, 14-3-3 (E), can first bind to either the MG (G) to form a binary complex (EG) or to the target protein (P) to form a different binary complex (EP). Subsequently, the final ternary complex (EGP) is formed when the EG complex binds with the target protein (P), or conversely, when the EP complex binds with the MG (G).

This system creates a non-dissipative thermodynamic cycle in which detailed balance must be satisfied; therefore, the dissociation constants are constrained by the relationship K_D1_K_D2_ = K_D3_K_D4_^13^. To impose this constraint, we define the parameter α such that K_D4_ = K_D1_/α and K_D2_ = K_D3_/α^14^. The parameter α represents the effective binding cooperativity due to the presence of the MG. A value of α>>1 indicates that the presence of MG has a strong cooperative effect on binding.

The equilibrium state of the system can then be described using three independent dissociation constants (*K*_*D*1_, *K*_*D*3_, and *α*) and the concentrations of the free species:

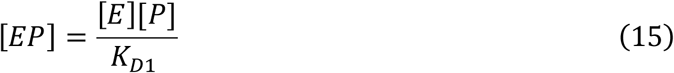

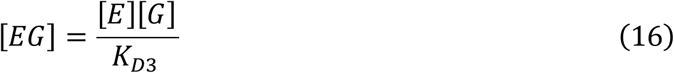

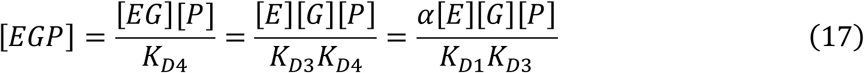

The total concentration of each component is conserved according to the mass balance equations:

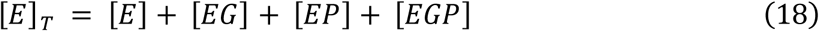

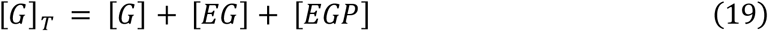

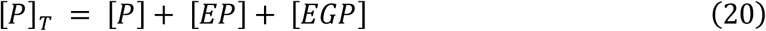

Solving this system of algebraic equations yields the steady-state concentration of each species as a function of the total MG concentration, *[G]_T_.* This allows us to model the signal from a nanoBRET assay, which measures the proximity between the effector protein (E) and POIs (P). The signal-generating species are the binary complex *[EP]* and the ternary complex *[EGP]*, as both involve the close association of E and P. To quantify the effect of the MG relative to the baseline interaction, the signal is normalized. Under the assumption that the BRET signal is linearly proportional to the total concentration of effector-target complexes (*[EP]+[EGP]*) the normalized nanoBRET signal (*X_BRET_*) is defined as the ratio of the total complex concentration at a given MG concentration (*[G]_T_*) to the concentration of the binary complex (*[EP]*) in the absence of any MG (*[G]_T_=0*). This relationship is expressed by the equation:

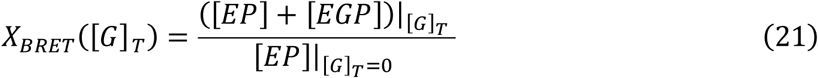

In this model, a value of *X_BRET_=1* represents the baseline interaction between the effector and target proteins. A value greater than 1 indicates that the MG enhances the association between the effector and target proteins, either by stabilizing the existing EP complex or by promoting the formation of the new EGP ternary complex.

### Mathematical Model of CRBN Degrader

The dynamics of CRBN MG degraders are described based on the identical binding network and cooperativity terms as the 14-3-3 MG stabilizers. However, unlike the stabilizer model which was analyzed at equilibrium, the degrader model must account for time-dependent protein dynamics. In this model, the POI concentration does not remain constant with time due to degradation. Therefore, an equilibrium assumption is no longer valid, and the system must be described dynamically. We therefore explicitly consider the forward and reverse rates for each of the four binding reactions in our model according to:

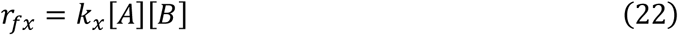

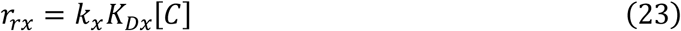

Where *r_fx_* the forward rate for binding reaction *x* (nM/hr), *r_rx_* is the reverse rate (nM/hr), *[A]* and *[B]* are the concentrations of the reactants (nM), *[C]* is the concentration of the binding product (nM), *k_x_*is the second-order association rate constant (1/nM/hr), and *K_Dx_*is the dissociation constant (nM). Terms describing the forward and reverse rates for each binding reaction are combined with terms representing baseline POI expression, degradation, and enhanced degradation through ternary complex formation to give the following six ordinary differential equations:

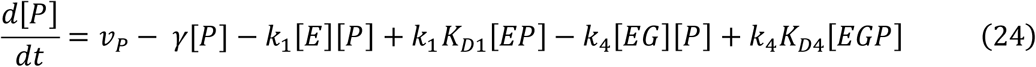

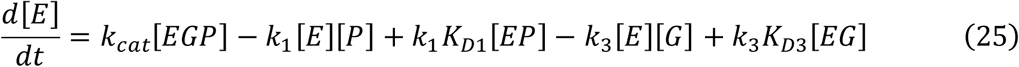

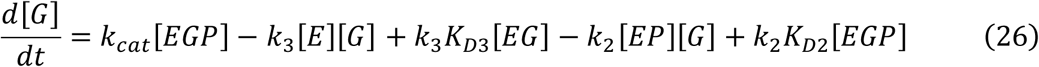

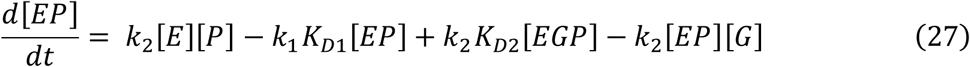

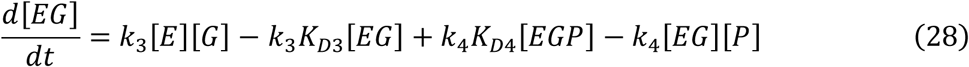

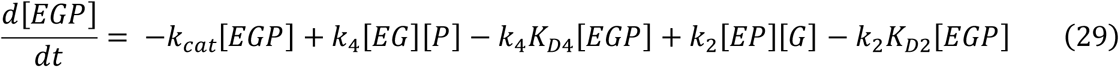

Where *[X]* is the concentration of species X (nM), *v_p_*is the baseline expression of the POI (nM/hr), *γ* is the baseline degradation rate constant of the POI (1/hr), and *k_cat_* is the rate constant of POI degradation from ternary complex (1/hr). Ubiquitination and degradation of the POI from ternary complex is represented as a single reaction that degrades the POI and recycles effector protein and MG to their free states. As a simplifying assumption, POI degradation from binary effector-POI (EP) complex was not considered. The parameters *K_D1_, K_D3_, α, k_cat_* are MG- and target-specific. For example, *k_cat_* for ternary complex between lenalidomide-CRBN-IKZF1 differs from that of lenalidomide-CRBN-GSPT.

The initial concentrations of non-MG species were calculated based on mass action equilibrium using the total amount of POI and effector protein, assuming the system is in a steady state prior to the addition of any MG. Specifically, the initial concentration for the POI (*[P]_0_*), binary effector-POI complex (*[EP]_0_*), and free effector protein (*[E]_0_)* are given by:

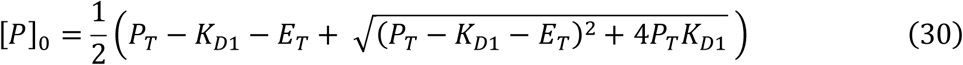

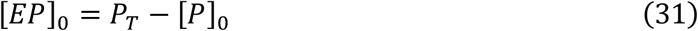

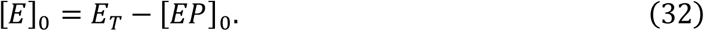

Where *P_T_* and *E_T_* are the total initial concentration of target and effector protein, respectively. The experimental condition simulated (i.e., iMRM or BRET) by this model determines the values of *E_T_* and *P_T_*. Furthermore, because iMRM experiments are conducted in wild-type cells, the value of *P_T_* depends on the POI being simulated. Because BRET experiments are conducted in engineered cells expressing engineered versions of both the POI and the effector protein (CRBN), we assume that the value of *P_T_* is constant for all POI in BRET experiments. Furthermore, we define baseline expression of the the POI, *v_p_*, such that the total POI and effector protein concentrations remain at *P_T_* and *E_T_,* according to:

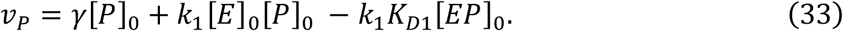

Finally, MG dosing is represented by setting the initial concentration of free MG (*[G]_0_*) in the system equal to the total concentration of MG defined by the dose according to:

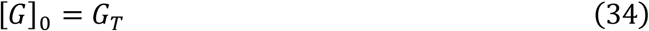

For simulating BRET^15^ or AlphaScreen^16^ experiments that measure ternary complex formation in absence of degradation, we set all degradation rate constants to zero. Furthermore, for fitting the combination of AlphaScreen and HiBiT data provided by Yamanaka and colleagues^16^, we assume that binding between CRBN and the POI or MG is weaker in the cell-free AlphaScreen assay because the additional species in the CRBN:DDB1:CUL4A:RBX1 that improve binding are not present^18,19^. Accordingly, we scale K_D1_ and K_D3_ for simulation of AlphaScreen experiments according to:

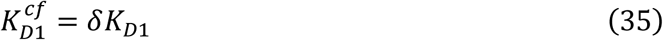

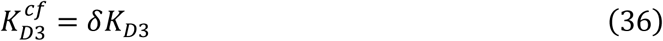

Where δ is the fitted scaling factor for cell free condition. The data describing MG degrader performance used for model validation comes in two forms: measurement of ternary complex formation in absence of degradation (BRET/AlphaScreen) and measurement of POI abundance (iMRM/HiBiT). Thus, two model different model outputs were used for data fitting depending on the experiment type. For fitting BRET and AlphaScreen data, we simulated the fold-increase in total target-effector (i.e., *[EP] + [EGP]*) complex after MG dosing compared to baseline according to:

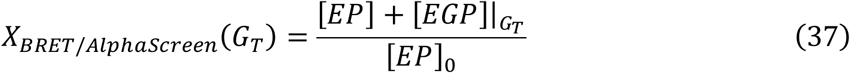

Similarly, for fitting iMRM and HiBiT data, we simulated the fraction of POI remaining after MG dosing compared to baseline according to:

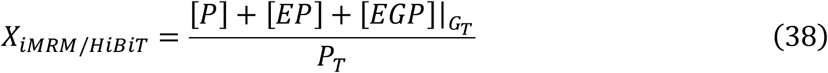

Note in our model, the total amount of POI at baseline is set by the parameter P_T_, so this value is set as the denominator in this equation. A complete list of model species and parameters are defined in **Table S3**.

### Parameter Estimation

Model parameters were estimated via an ensemble approach using global optimization to obtain a large number of parameter sets with good fitness to data^20,21^. Models were fitted independently to datasets from each publication by minimizing the weighted sum of squared errors (SSE) between model predictions and experimental measurements across all MG:POI combinations:

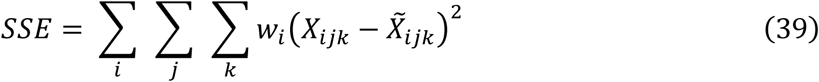

where *w_i_* represents the weight for experiment type *i* (e.g., BRET), *X_ijk_* is the model prediction, and 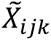 is the measured value for experiment type *i* and MG: POI combination *j*, *k*. Most experiments were assigned a weight of 1, with two exceptions: covalent modification data from the CYPA system by Schulze et al. (w = 10,000) and the HiBiT dataset from Yamanaka et al. (w = 100). These weights were selected to ensure that different experiment types contributed approximately equally during fitting.

Model parameters were estimated by fitting the combined dataset from each literature source separately. Global optimization was implemented using MATLAB’s particleswarm algorithm with a different random seed each run, and an ensemble of parameter sets within 5% of the lowest value were selected for analysis. Specifically for the MG stabilizer, the selection was restricted to conditions where the total effector protein concentration exceeded the total POI concentration (i.e., [*E*]_*T*_ > [*P*]_*T*_), as its stabilizing effect is valid only in this regime. Optimization was implemented such that at least 10 parameter sets were generated for each literature source.

### Sensitivity Analyses

Sensitivity analyses were implemented by calculating the log-gain sensitivity of predicted MG behavior to each model parameter according to:

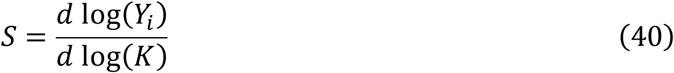

Where *Y_i_* is the predicted measure of MG performance (D_max_, TC_max_, EC_50_) K and is the model parameter. Sensitivities were calculated numerically by simulating our models before and after a small change in the parameter value (1%).

## Results

### The mathematical model framework accurately represents data across a range of molecular glue contexts

To systematically investigate the relationship between MG kinetic parameters and performance, we developed mathematical models for MG stabilizer and MG degrader modalities (**Figure 1, Supplementary Tables S1-4**). Given the mechanistic diversity among MGs, we constructed a suite of models to capture key variations in their modes of action. These include: (i) a minimal single-pathway ternary complex formation model (**Figure 1A**), (ii) a model incorporating covalent modification of ternary complex (**Figure 1B**), (iii) a two-pathway ternary complex formation model (**Figure 1C**), and (iv) a model representing ternary complex-mediated POI degradation (**Figure 1D**). Across all models, G represents the MG, P represents the POI, and E represents the effector protein. The specific proteins and molecules corresponding to the E, G, and P components in each model are listed in **Supplementary Table S4**.

**Figure 1.**
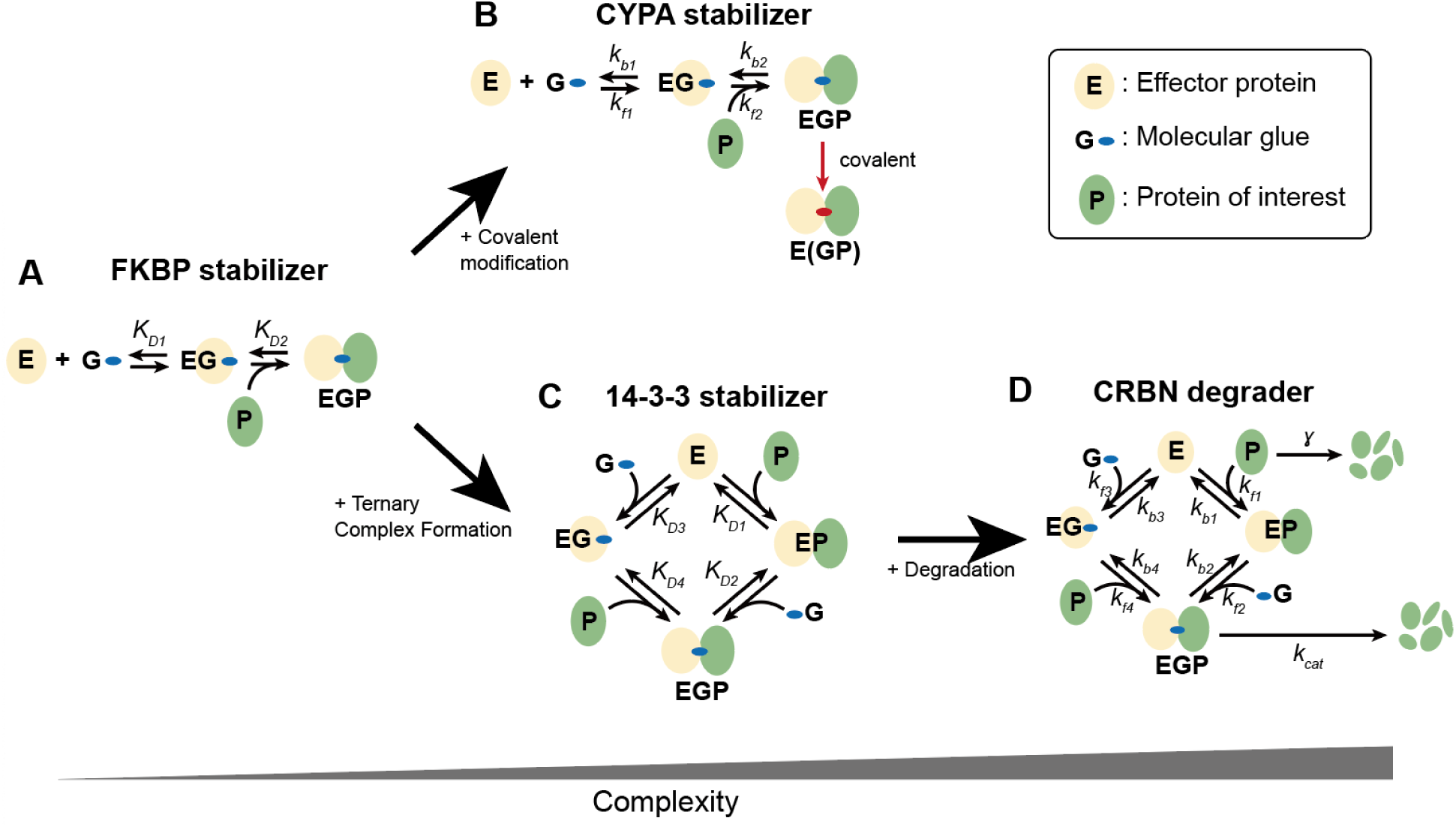
Unified kinetic framework to understand MG mechanisms with increasing complexity. Models describing MG dynamics of (A) the FKBP stabilizer, representing a minimal single-pathway model; (B) the CYPA stabilizer, which adds a covalent step; (C) the 14-3-3 stabilizer, which introduces a second pathway for complex formation; and (D) the CRBN degrader, which includes the final step of protein degradation. E: effector protein. P: POI. G: MG.

We validated these models by fitting model predictions to multiple datasets compiled from published studies, comprising measurements obtained via Fluorescence Polarization (FP)^22^, Bioluminescence Resonance Energy Transfer (BRET)^23^, Time-Resolved Fluorescence Resonance Energy Transfer (TR-FRET)^24^, Nano-Glo HiBiT Lytic Detection (HiBiT)^17^, AlphaScreen (AS)^16^, and immuno-Multiple Reaction Monitoring (iMRM) assays^15^ (**Table S3**). To assess parameter identifiability, parameter optimization was implemented using an ensemble approach, in which many well-fitting parameter sets are obtained and evaluated. MG stabilizer models were validated based on their ability to replicate FP (**Figure 2A**), TR-FRET (**Figure 2B**), covalent modification (**Figure 2C**), and nanoBRET (**Figure 2D, Figure S1**) datasets, and MG degrader models based on BRET (**Figure 3A, Figure S2**), iMRM (**Figure 3B-C, Figure S3-4**), AlphaScreen (**Figure 3D, Figure S5A-C**), and HiBiT (**Figure 3E, Figure S5D-F**) datasets. Both the MG stabilizer and degrader models effectively recapitulated the experimental data, demonstrating strong overall agreement.

**Figure 2:**
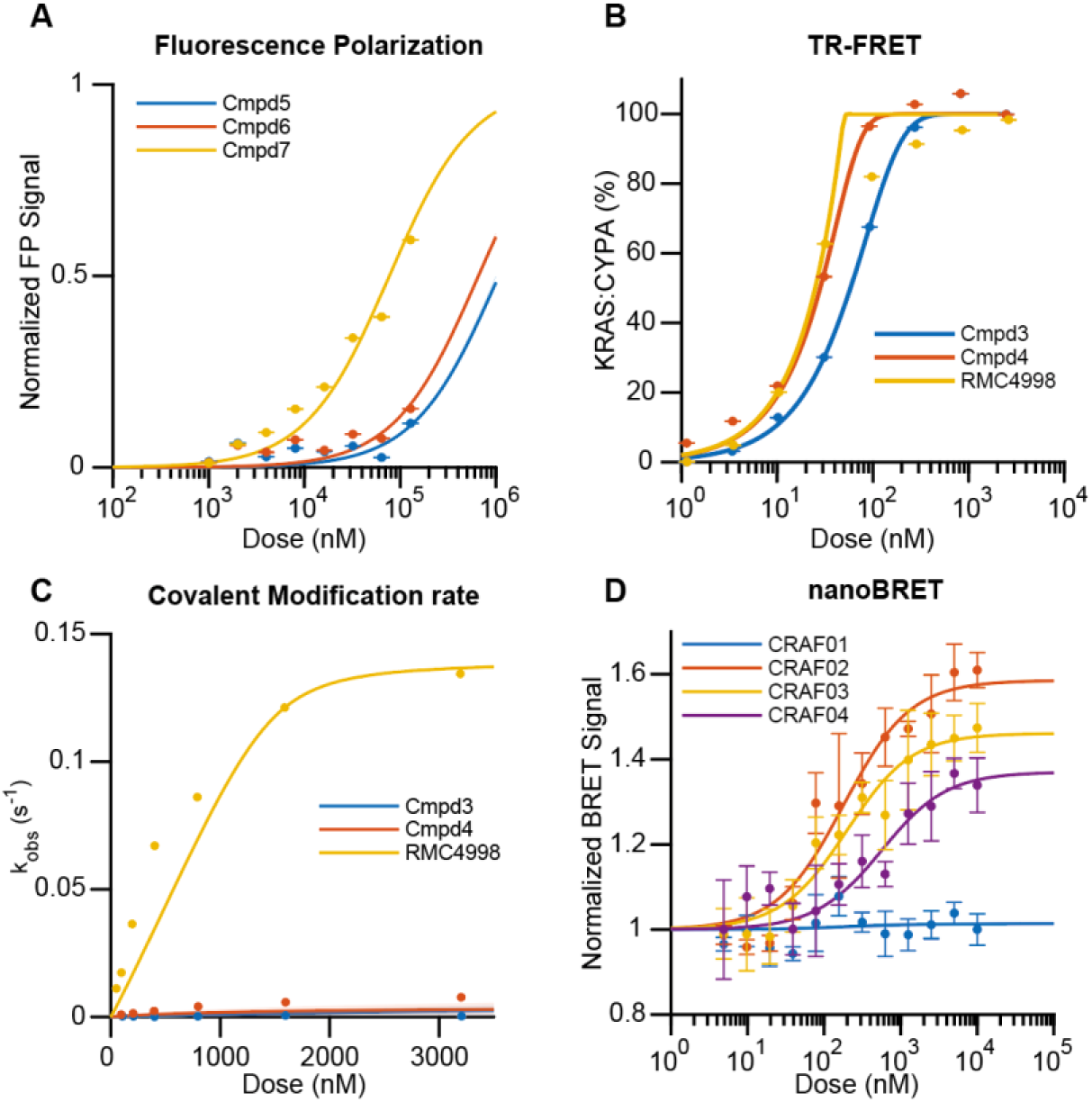
Fitted MG stabilizer model predictions compared to experimental data. (A) Ensemble (n=30) model fitness, based on the model in Figure 1A, to FP dose–response data from Deutcher et al., describing ternary complex (FKBP12:MG:FRB) formation induced by Cmpd5 (blue), Cmpd6 (red), and Cmpd7 (yellow). (B-C) Ensemble (n=30) model fitness based on the model in Figure 1B, to (B) TR-FRET dose–response data and (C) covalent modification rate data from Schulze et al., describing ternary complex (CYPA:MG:KRAS) formation induced by Cmpd3 (blue), Cmpd4 (red), and RMC4998 (yellow). (D) Ensemble (n=24) model fitness, based on the model in Figure 1C, to nanoBRET data from Vickery et al., describing ternary complex (14-3-3:MG:CRAF) formation induced by CRAF01 (blue), CRAF02 (red), CRAF03 (yellow), and CRAF04 (purple). Solid curves: Mean predicted values. Shaded regions: Raw 95% interval. Points: experimental data.

**Figure 3:**
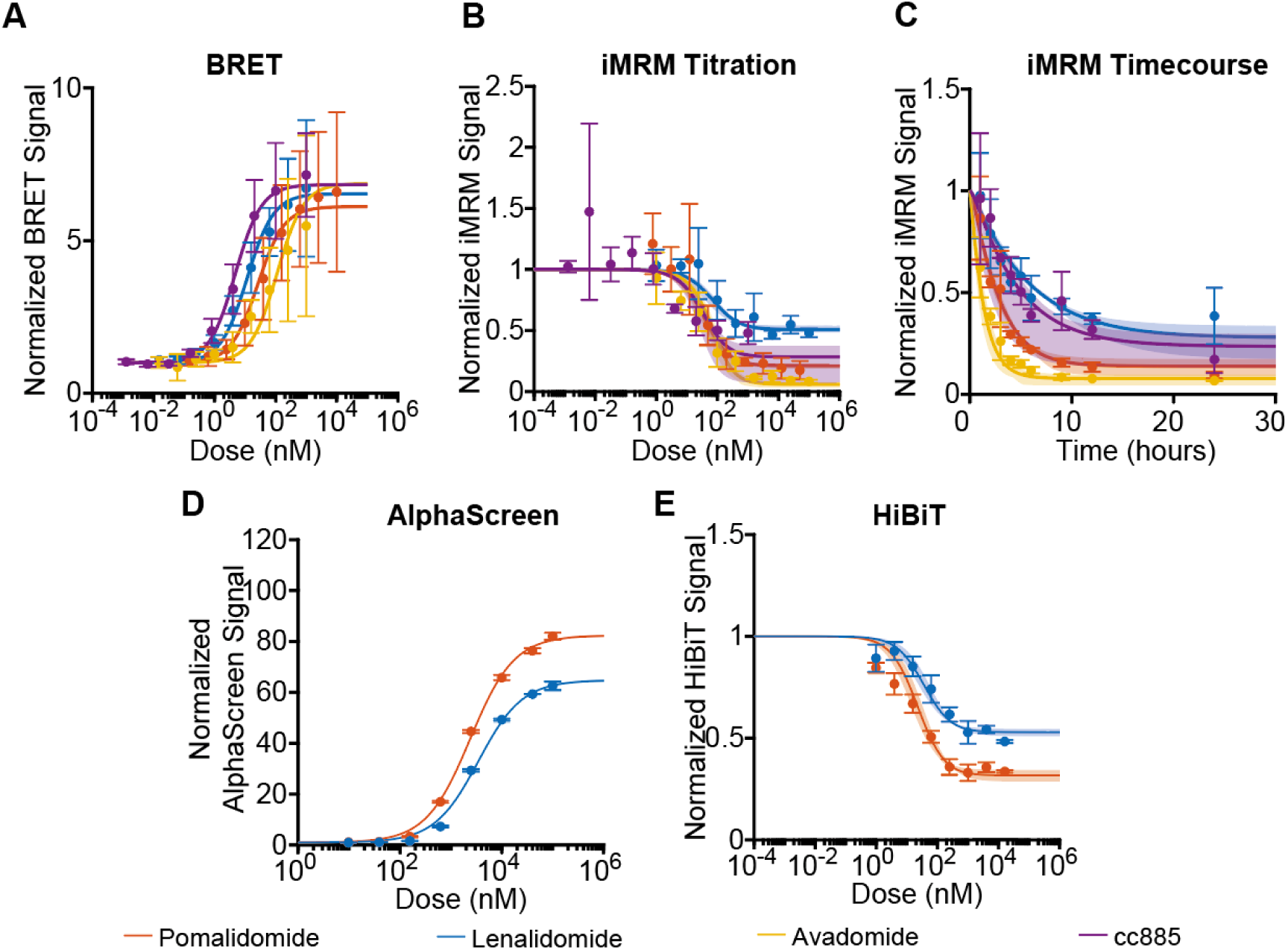
Fitted MG degrader model predictions compared to multiple experimental datasets. (A-C) Ensemble (n = 29) model fitness to (A) BRET data, (B) iMRM dose response data, and (C) iMRM timecourse data provided by Sperling et al. describing pomalidomide (orange), lenalidomide (blue), avadomide (yellow), and cc885 (purple) binding to and induced degradation of IKZF1. (D-E) Ensemble (n = 10) model fitness to (D) AlphaScreen data and (E) HiBiT data provided by Yamanaka et al., describing pomalidomide (orange) and lenalidomide (blue) binding to and induced degradation of IKZF1. All fits generated using model depicted in Figure 1D. Solid curves: Mean predicted values. Shaded regions: Raw 95% interval. Points: experimental data.

The estimated parameters from model fitting provided mechanistic insights into determinants of MG performance. Notably, our models suggest that more effective stabilizers exhibit higher MG-mediated cooperativity, while more effective degraders are characterized by stronger binding and higher catalytic efficiency. The ensemble modeling revealed that some parameters remained loosely constrained (**Supplementary Tables S5-9**), likely due to limitations of normalized measurements, which restricted identifiability of absolute kinetic rates. Nevertheless, the overall model fits were consistent, indicating that our framework successfully captures the essential dynamics of MG-induced interactions and downstream effects.

### Affinity of the E:MG complex for the POI controls performance across all MG modalities under fixed stoichiometry and degradation rates

Using the validated models, we systematically evaluated how variations in kinetic parameters affect the performance of each MG modality. To quantify performance, we simulated dose-response curves by varying the MG concentration for any given parameter set and extracted two key metrics: maximal engagement and potency (EC_50_) (**Figure 4**). Maximal engagement was defined as the maximum fraction of the POI forming a ternary complex (TC_max_) for stabilizers (**Figure 4A**), or the maximum fraction degraded (D_max_) for degraders (**Figure 4B**) at saturating MG concentrations. In both cases, potency was defined as the concentration of MG required to reach 50% of the maximal engagement.

**Figure 4:**
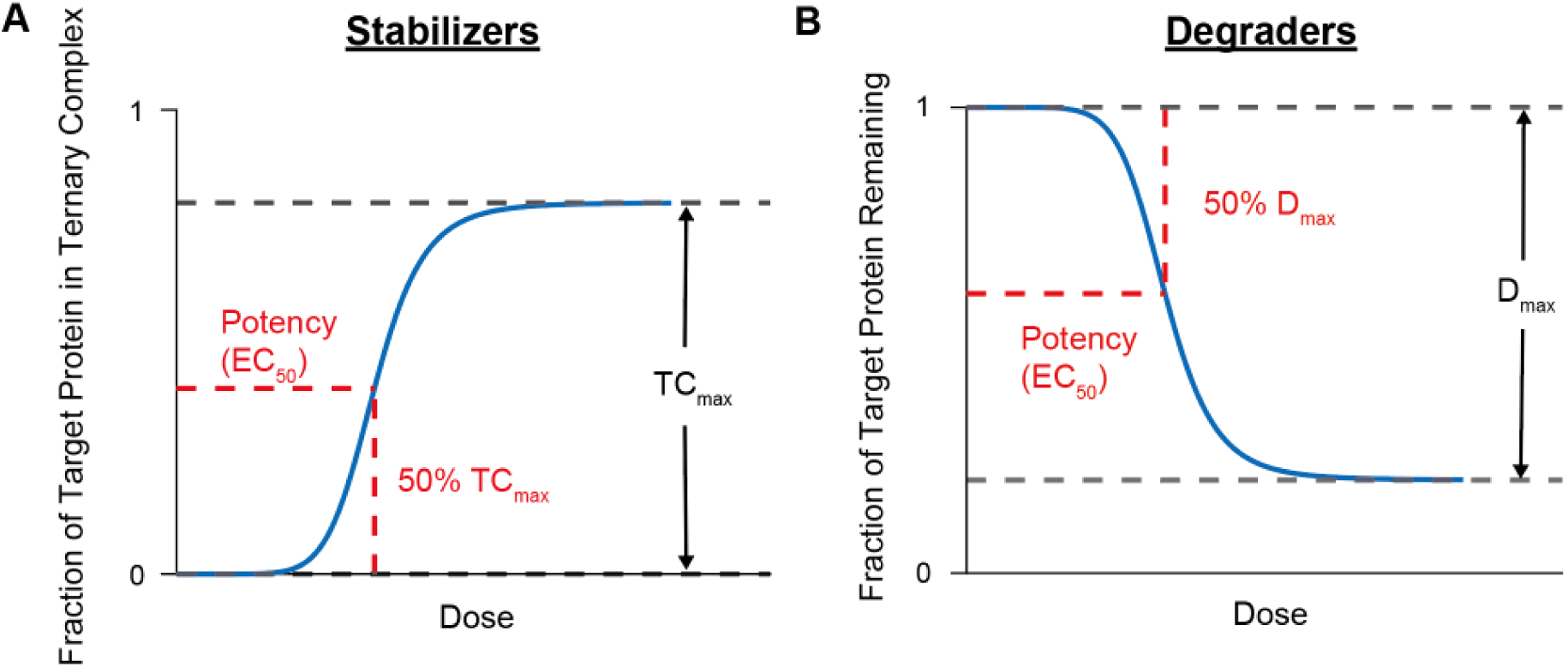
Metrics for quantifying MG performance. (A) The performance of MG stabilizer is assessed by the maximal ternary complex formation (*TC*_*max*_) and potency (*EC*_50_). (B) The performance of MG degrader is assessed by the maximal target protein degradation (*D*_*max*_) and potency (*EC*_50_).

We first investigated the effects of binding affinity and cooperativity on the predicted performance of MG stabilizers, utilizing our validated 14-3-3 model (**Figure 1C**). This analysis utilized the best-fit parameter set for the CRAF02 stabilizer, derived from the CRAF dataset (**Supplementary Table S10**)^8^. We then systematically calculated the maximal ternary complex formation (TC_max_) and potency (EC_50_) while varying the binding affinities (K_D1_, K_D3_) and cooperativity (α). Remarkably, across all parameter combinations, the TC_max_ was determined exclusively by the value of K_D4_, the dissociation constant of the POI from the MG:effector protein complex (**Figure 5A**). This suggests that under saturating MG conditions, the system is primarily governed by the equilibrium between the EG and EGP forms, making the stability of the final ternary complex the sole determinant of maximal effects. Supporting this, variations in K_D3_ (E:G affinity) had no impact on TC_max_ (**Figure 5B**). Changes in cooperativity (α) affected TC_max_ only through their effect on K_D4_ (since K_D4_ = K_D1_/α). In contrast, stabilizer potency (EC_50_) improved as binding strength increased for all parameters (**Figure 5C-D**). This indicates that while maximal engagement is dependent only on the final complex stability, the efficiency of reaching that state—the potency—depends on the broader kinetics of all binding events.

**Figure 5:**
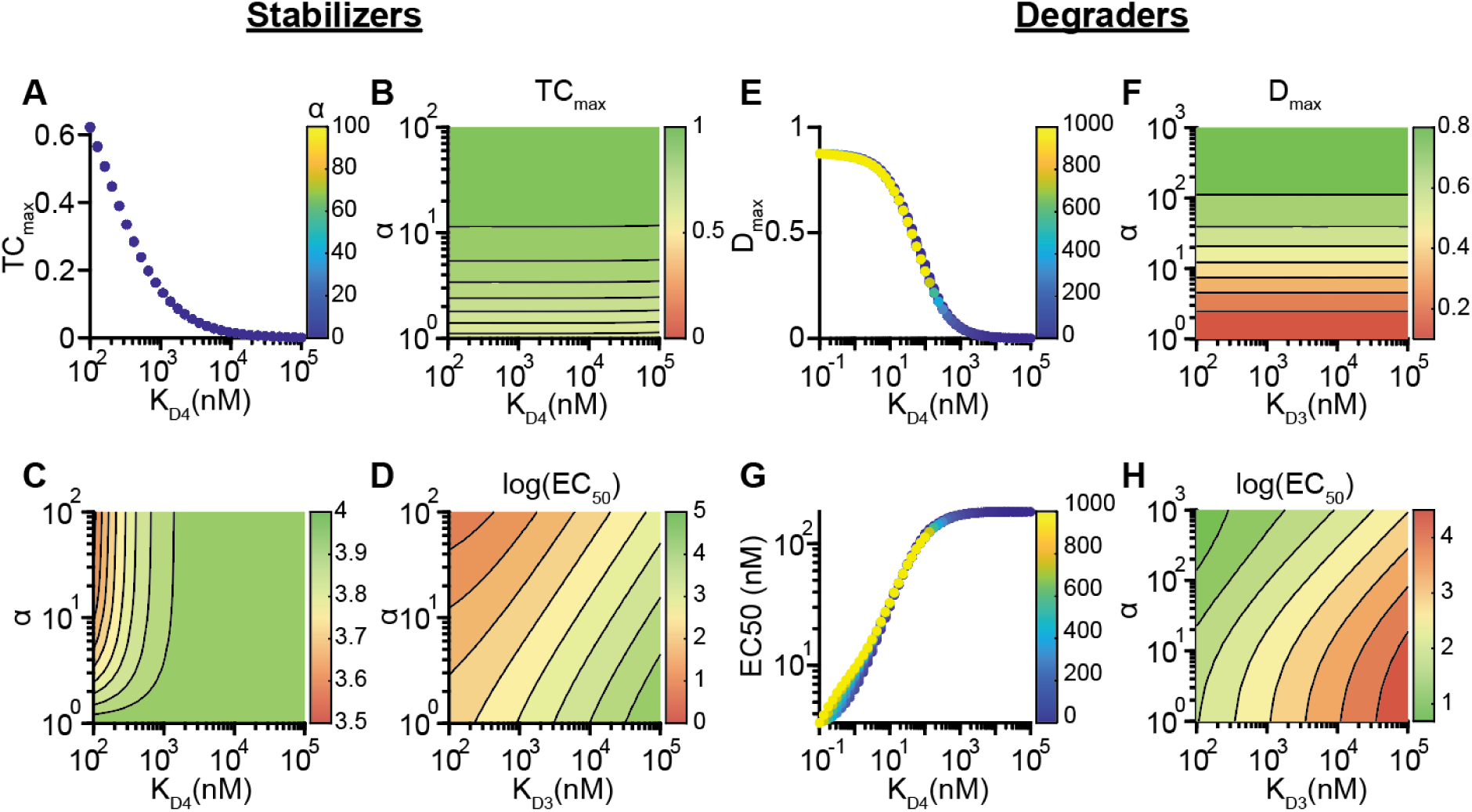
Predicted effects of MG binding affinities on drug performance. (A) Predicted value of *TC*_*max*_ for CRAF02-induced stabilization of CRAF varying *K*_*D*4_. (B) Heatmap illustrating predicted effects of cooperativity and baseline E:G binding on predicted *TC*_*max*_. (C-D) Heatmaps illustrating predicted effects of cooperativity and EG:P binding (C) and cooperativity and baseline E:G binding on predicted EC_50_ (E) Predicted value of *D*_*max*_ for avadomide-induced IKZF1 degradation varying *K*_*D*4_. (F) Heatmaps illustrating predicted effects of cooperativity and baseline E:G binding on predicted ^*D*^*max*. (G) Predicted value of EC_50_ for avadomide-induced IKZF1 degradation varying *K*_*D*4_. (F) Heatmaps illustrating predicted effects of cooperativity and baseline E:G binding on predicted EC_50_.

We next analyzed the effects of binding affinity and cooperativity on the predicted performance of MG degraders. For these analyses, we simulated the lenalidomide-mediated degradation of IKZF1 using the best-fit parameter set to the data presented by Sperling et al. (**Supplementary Table S11**)^15^. To assess the effects of binding affinities, we calculated D_max_ and EC_50_ by varying MG-induced cooperativity (α) simultaneously with either E:P binding affinity (K_D1_) or E:G binding affinity (K_D3_), with all other parameters (i.e., species abundances and degradation rate constant) fixed. Like stabilizers, the maximum effect of degraders depended solely on K_D4_. Simulated D_max_ values, obtained by varying α and K_D1_ simultaneously, collapsed onto a single curve described by K_D4_ = K_D1_/α (**Figure 5E**), and we found that K_D3_ did not affect D_max_ (**Figure 5F**). In contrast, all binding parameters influenced the predicted EC_50_. Specifically, while the simulated EC_50_ values varying α and K_D1_ similarly collapse onto a single curve with slight dependencies on α at low K_D4_ (**Figure 5G**), varying α and K_D3_ also strongly influenced EC_50_ (**Figure 5H**). Notably, the EC_50_ values obtained by covarying K_D3_ and α span a larger range than those obtained by varying K_D1_ and α simultaneously, suggesting that K_D3_ more strongly influences EC_50_. Overall, our findings demonstrate that MG stabilizers and degraders exhibit similar parameter-performance relationships. Maximum engagement (D_max_ and TC_max_) is solely determined by ternary complex binding affinity K_D4_, whereas potency (EC_50_) improves with stronger binding for all key parameters.

### In the limit of very strong binding, MG degraders are limited by catalytic efficiency

MG degrader performance should also be affected by the catalytic rates of POI degradation, especially when the binding is strong. To analyze this effect, we first simulated the effects of varying k_cat_, the effective target protein degradation rate constant from the MG-induced ternary complex, and K_D4_ as the limit of irreversible binding is approached (K_D4_ → 0). Similar to our results in the previous section, stronger ternary complex binding (i.e., decreasing K_D4_) leads to greater achievable POI degradation; however, when binding becomes sufficiently strong D_max_ depends solely on k_cat_ (**Figure 6A**). The predicted EC_50_ values also decrease as K_D4_ decreases until a threshold is reached, below which k_cat_ controls EC_50_ (**Figure 6B**). Ultimately, these results demonstrate that in the limit of high binding affinity, the overall performance in terms of both D_max_ and EC_50_ of MG degraders is determined by the catalytic efficiency of the ternary complex.

**Figure 6:**
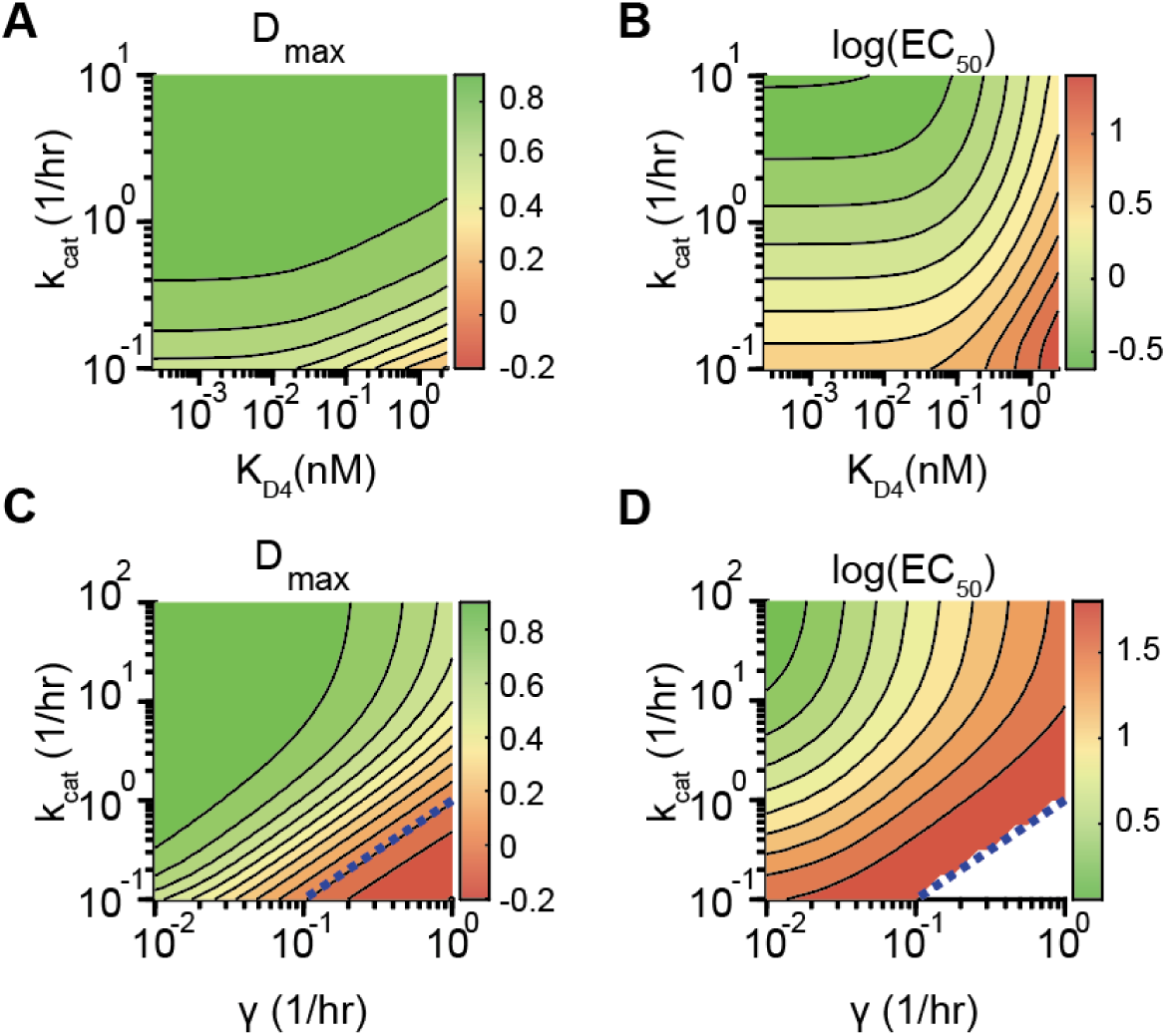
Effective rate of MG-induced degradation limits MG degrader performance. (A-B) Predicted effects of co-varying *k*_*cat*_ and *K*_*D*4_ on A) *D*_*max*_ and B) EC_50_. (C-D) Predicted effects of co-varying *k*_*cat*_ and *γ* on C) *D*_*max*_ and D) EC_50._ White region in panel D) indicates no EC_50_ was calculated because *D*_*max*_ is negative.

We expect the baseline degradation rate of the POI rates in the absence of MG to influence MG performance. To quantify this, we analyzed the effects of covarying k_cat_ and γ, the rate constant governing baseline POI degradation. Our model predicts that D_max_ inversely correlates with γ, but significant POI degradation is achievable if k_cat_ is sufficiently high (**Figure 6C**). Interestingly, a transition from positive to negative D_max_ occurs at the y = x line, demonstrating that extreme cases where POI degradation from the ternary complex is reduced compared to the natural degradation can lead to increased POI abundance. Similarly, potency decreases with higher γ, but higher k_cat_ values can compensate and restore potency (**Figure 6D**). These results together indicate that to effectively degrade a POI with a high D_max_, the effective degradation rate constant (k_cat_) from the MG-mediated ternary complex must be sufficiently greater than the natural POI degradation rate constant (γ). Consequently, it is more challenging to develop a MG degrader against a POI with higher natural degradation rate constant.

### Abundance of the effector protein limits performance in both MG stabilizers and degraders

While kinetic parameters are engineerable properties of an MG, the cellular environment, particularly protein abundance, can profoundly influence its performance. Therefore, we next investigated the impact of protein abundance levels on the performance of MG stabilizers. Using the 14-3-3 model, we first examined how TC_max_ and EC_50_ change in response to varying both POI and effector protein concentrations. Our model predicts that TC_max_ increases with higher effector protein concentration and lower POI concentration (**Figure 7A**). Notably, when the concentration of the POI is significantly lower than that of the effector protein, the system is less sensitive to changes in POI abundance. However, TC_max_ decreased sharply as the stoichiometry of the two proteins deviated from a 1:1 ratio. This is expected, as MG stabilizers require one effector protein to stabilize one POI, highlighting that stoichiometry is a critical determinant of a stabilizer’s maximal effect. Regarding potency, we found that the EC_50_ decreased (indicating higher potency) as the concentration of either the effector or the target protein increased (**Figure 7B**). For a stabilizer to be effective, it is therefore essential that the effector protein concentration exceeds that of the POI.

**Figure 7:**
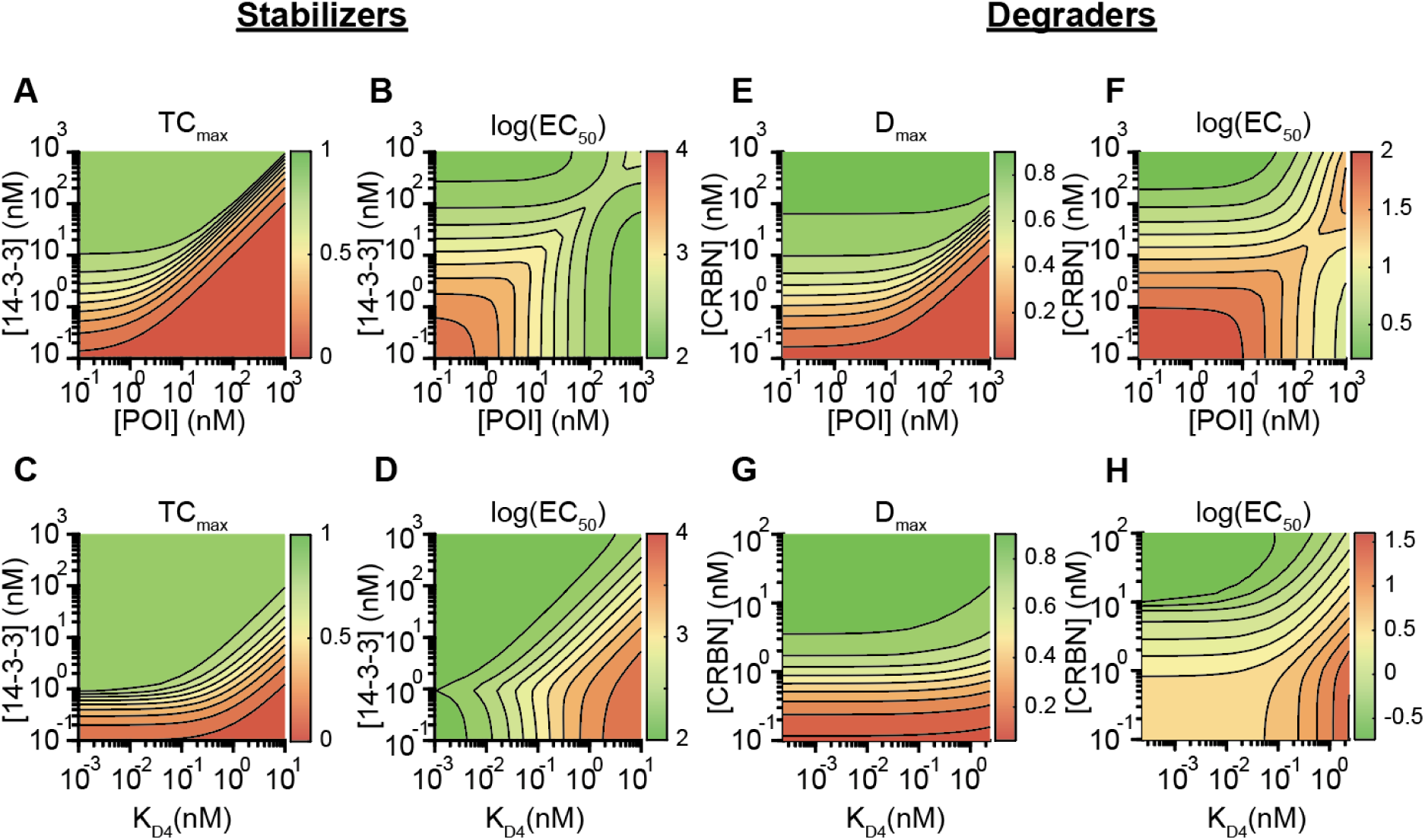
Predicted effects of target and effector protein expression on MG performance. (A-B) Heatmaps demonstrating the effects of target protein expression (x-axis) and effector protein expression (y-axis) on (A) *TC*_*max*_ and (B) EC_50_ for CRAF02-induced stabilization of CRAF. (C-D) Heatmaps demonstrating the effects of EG:P binding affinity(x-axis) and effector protein expression (y-axis) on (C) *TC*_*max*_ and (D) EC_50_ for CRAF02-induced stabilization of CRAF. (E-F) Heatmaps demonstrating the effects of target protein expression (x-axis) and effector protein expression (y-axis) on (E) *D*_*max*_ and (F) EC_50_ for lenalidomide-induced degradation of IKZF1. (G-H) Heatmaps demonstrating the effects of target protein expression (x-axis) and EG:P binding affinity (y-axis) on (G) *D*_*max*_ and (H) EC_50_ for lenalidomide-induced degradation of IKZF1.

To explore whether a strong binding affinity could compensate for the scarcity of the effector protein, we extended our analysis by varying the effector protein concentration and the ternary complex stability (K_D4_). We observed that once the binding affinity becomes sufficiently strong (i.e., below a certain K_D4_ threshold), further increases in affinity no longer improve TC_max_ (**Figure 7C**). In this regime, the maximal effect becomes solely limited by the effector protein concentration. This analysis reinforces that effector protein abundance and the resulting stoichiometry are crucial factors governing the performance of MG stabilizers (**Figure 7D**).

We next investigated the effects of POI and effector protein abundance on the predicted performance of MG degraders. Like MG stabilizers, our model for MG degraders predicts poor degrader performance (D_max_) when the POI becomes sufficiently high (**Figure 7E-F**). However, in contrast to stabilizers, there is a larger region where the POI concentration has minimal effect on performance. These results indicate that MG degraders perform effectively over a broader range of effector protein: POI ratios. Conversely, higher effector protein concentrations uniformly lead to higher D_max_ and improved EC_50_. These results ultimately suggest that MG degraders are relatively insensitive to target POI abundances, but the concentration of effector protein is a critical parameter. To explore this further, we analyzed the effects of covarying effector protein concentration and K_D4_ in the regime of strong binding. Like our previous results, lower K_D4_ values are associated with improved D_max_ (**Figure 7G**) and EC_50_ (**Figure 7H**), but the system ultimately becomes limited by effector protein concentration when binding is sufficiently strong. Thus, the concentration of effector protein is another system-determining parameter in the limit of optimal binding affinities.

### Sensitivity analyses illustrate that the predicted effects of model parameters are conserved across optimized parameter sets

To ensure that the uncertainty in the estimated model parameters does not alter our conclusions, we implemented local sensitivity analyses. More specifically, we calculated the log-gain sensitivities of MG performance characteristics (TC_max_, D_max_, and EC_50_) to each model parameter for every parameter set in the ensemble obtained from data fitting (**Figure 8**). The same MG:POI combinations were used here as in the previous sections. Interestingly, the sensitivities of EC_50_ to model parameters were larger in magnitude compared to D_max_ for MG degraders (**Figure 8C-D**). Furthermore, our analysis shows that despite the wide variance in some estimated parameters, the model’s performance sensitivities remain consistent. Specifically, the sensitivities of MG performance characteristics to any given parameter never changed sign, and their magnitudes remained stable across the parameters’ respective uncertainty ranges, with no extreme variations observed. Altogether, this demonstrates that our conclusions are robust and independent of the specific parameter values used in the simulation.

**Figure 8:**
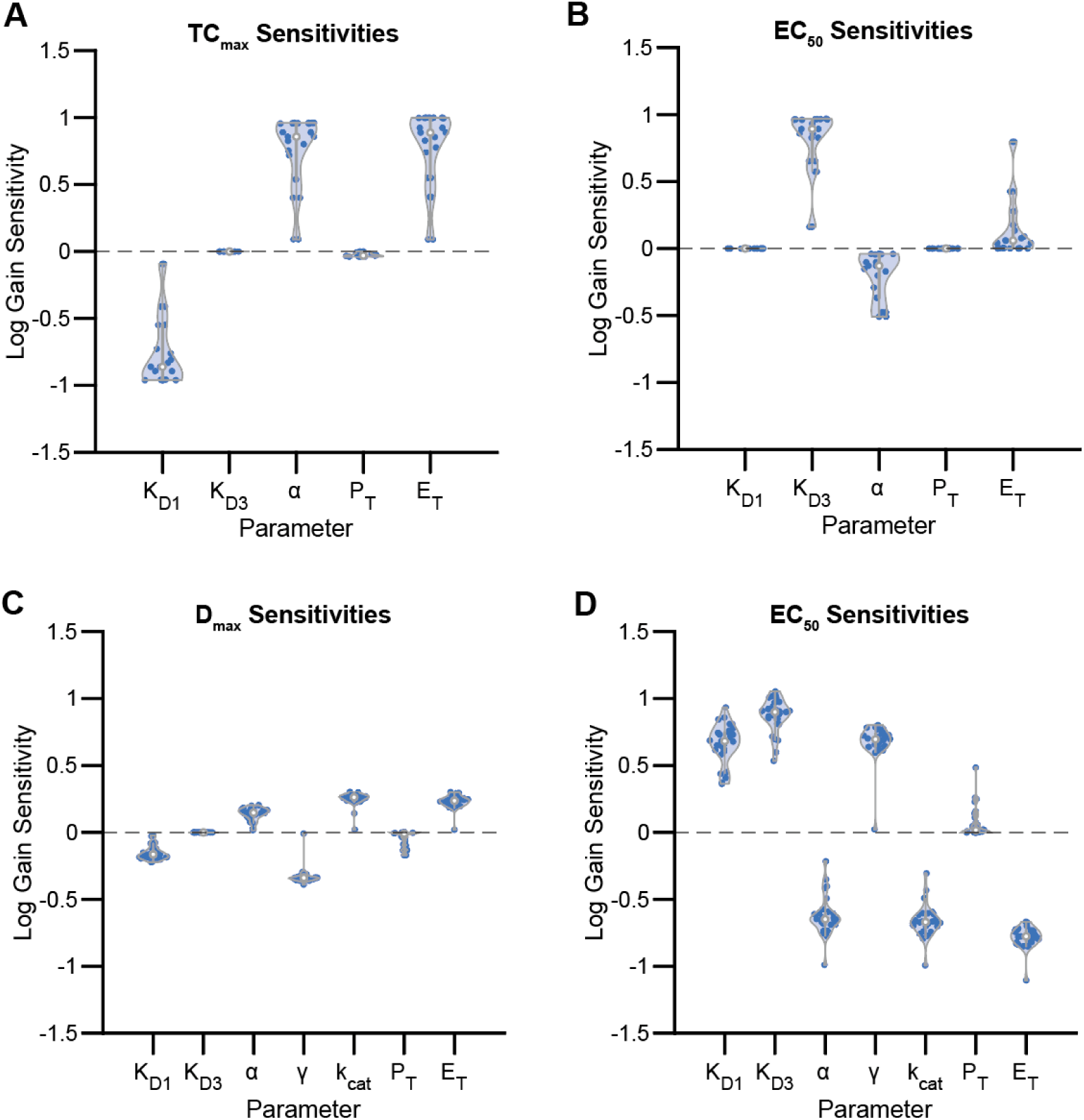
Calculated log-gain sensitivities of model parameters to MG performance characteristics. (A-B) Log gain sensitivity of A) TC_max_ and B) EC_50_ to MG stabilizer model parameters. (C-D) Log gain sensitivity of C) D_max_ and D) EC_50_ to MG degrader model parameters. Plots generated using code provided by Bastian Bechtold (https://github.com/bastibe/Violinplot-Matlab).

## Discussion

In this work, we developed a unified kinetic framework to describe the mechanisms of action for both MG stabilizers and degraders. We demonstrated that these models contain the necessary features to capture experimental trends across multiple diverse datasets accurately. By systematically analyzing this framework, we have derived a set of mechanistic principles that provide guidelines for the rational design and successful development of MGs.

Our findings provide key insight into the relationship between MG parameters and performance. We categorize these parameters into two distinct groups: MG intrinsic parameters and extrinsic parameters. MG intrinsic parameters are parameters that can be optimized through the chemical modification of MG molecules, i.e., MG:effector protein binding affinity and cooperativity. In contrast, extrinsic parameters cannot be directly optimized but must be addressed through rational selection of effector proteins and POIs. Parameters in this category include baseline effector:POI binding, baseline POI degradation rate, and expression of both proteins. We also classify the effective POI degradation rate from the ternary complex as extrinsic because it is ultimately limited by the maximum degradation capacity of the ubiquitin-proteosome system, which cannot be modified through optimization of the MG molecule.

For both MG stabilizers and degraders, the ternary complex binding affinity K_D4_, the dissociation constant of POI binding to the MG:effector protein complex, determines the maximum effect in terms of TC_max_ or D_max_, but higher cooperativity and stronger MG:effector protein binding also increase potency. However, strong binding alone is insufficient to maximize the MG performance, and the extrinsic parameters of MG ultimately limit the system. For MG stabilizers, a high relative expression of the effector protein compared to the POI is required for sufficient inhibition. Even with ideal, near-irreversible binding, significant POI inhibition is impossible if this condition is not satisfied. Accordingly, the selection of POIs and effector proteins is crucial for designing MG stabilizers. Effector proteins with high levels of expression in target tissue must be selected, such as 14-3-3σ, the effector protein for many MG stabilizers^9^. Similarly, MG degrader performance is ultimately limited by the effective POI degradation rate from ternary complex and the expression of the effector protein. Therefore, effector protein selection remains important for degraders; however, ensuring that the ternary complex is catalytically active in early-stage development is also critical. More specifically, the formation of a ternary complex must lead to effective POI degradation sufficiently faster than the baseline degradation rate. MG degraders have been shown in the literature to exhibit distinct degradation kinetics despite having similar binding^15^, supporting the notion that these parameters are critical for MG degrader development. Overall, our findings reveal that the successful development of an MG requires strong consideration of these extrinsic properties, in addition to optimizing binding and cooperativity.

This study has several limitations. First, our models were validated with published datasets using assays that primarily do not report relative abundances (**Figure S4**), which challenges the precise identification of absolute kinetic parameters. However, our sensitivity analyses demonstrate that the study’s major conclusions are robust and remain consistent across a wide range of parameter values (**Figures S6-7**). Second, a lack of detailed data on protein modifications, such as ubiquitination or other covalent modifications, necessitated a simplified model structure. For instance, while in the reported example, 14-3-3 σ protein undergoes a covalent modification step, our model simplifies this interaction to reversible binding and unbinding due to the lack of reported kinetic data on the covalent step. Although our current conclusions are robust, future work incorporating more comprehensive data could refine the model to capture these complex mechanisms more realistically.

There are many MG degraders and stabilizers in both clinical and preclinical development today. Most of these molecules are designed for the treatment of hematological^25–32^ and solid^33–35^ tumors; however, MGs are also being developed for other diseases such as systemic lupus erythematosus^36^ and colitis^37^. Due to the variety of downstream effects CIP can induce, including both upregulation and downregulation of disease-relevant pathways, MGs can treat a wide range of conditions as development continues. However, MG development has been complicated by the emergence of resistance^38^ or difficulty identifying clear prognostic markers^39,40^. Cell-level features, such as expression of target and effector proteins, are central to these challenges, and our kinetic framework elucidates the effects of these parameters. Furthermore, our model offers a direct path to accelerating MG development. By incorporating measurable parameters—dissociation constants, protein expression levels, and degradation rates—the model enables *in silico* predictions that can streamline early-stage go/no-go decisions during risk assessment of new MG projects, to highlight key data needed to be generated to understand MG behavior, and ultimately to guide specific MG optimization strategies. A particularly promising extension would integrate this cellular kinetic model with whole-body pharmacokinetic and pharmacodynamic models. Such integration would illuminate how molecular-level parameters translate to clinically relevant properties, including plasma half-life, tissue distribution, and therapeutic window, ultimately bridging the gap between biochemical mechanisms and therapeutic outcomes.

## Supporting Information

Supplementary tables, figures, and text are located in the attached file “Supplementary Information.pdf”. All code used for model simulation and parameter estimation is found at the following GitHub repository: https://github.com/jcd111/MG_Manuscript_Model.

## Acknowledgements

This work was funded by AstraZeneca and collaboration support provided by the AstraZeneca Open Innovation program. The simulations were performed on the Rice University’s Center for Research Computing equipment acquired with the Big-Data Private-Cloud Research Cyberinfrastructure MRI-award funded by the NSF under grant CNS-1338099. We would also like to thank Dr. Adam Sperling et al. and Dr. Michelle Arkin et al. for providing raw data for model validation.

## SUPPLEMENTARY INFORMATION

## Supplementary Tables

**Supplementary Table S1:** Description of MG stabilizer model parameters for FKBP/CYPA systems.

| Parameter | Meaning | Source |
| --- | --- | --- |
| $K_{D1}$ | Baseline E:G binding affinity. Different for every E:G combination. | Fixed to reported values. |
| $K_{D2}$ | Binding affinity between EG binary complex and P. Different for every EG:P combination. | Estimated by fitting to data. |
| $P_T$ | Abundance of target protein. Different for each assay. | Fixed to reported values. |
| $E_T$ | Abundance of effector protein. Different for each assay. | Fixed to reported values. |
| $k_{cat}$ | Covalent modification rate of EGP ternary complex (CYPA model only) | Estimated by fitting to data. |

**Supplementary Table S2:** Description of MG stabilizer model parameters for 14-3-3 system.

| Parameter | Meaning | Source |
| --- | --- | --- |
| $K_{D1}$ | Baseline E:P binding affinity. Different for every E:P combination. | Estimated by fitting to data. |
| $K_{D3}$ | Baseline E:G binding. Different for every E:G combination. | Fixed to either reported or approximated values. |
| $\alpha$ | Cooperativity of MG binding. | Estimated by fitting to data. |
| $P_T$ | Abundance of target protein. Different for every E:P combination. | Estimated by fitting to data. |
| $E_T$ | Abundance of effector protein. Different for every E:P combination. | Estimated by fitting to data. |

**Supplementary Table S3:** Description of MG degrader model parameters.

| Parameter | Meaning | Source |
| --- | --- | --- |
| $K_{D1}$ | Baseline E:P binding affinity. Different for every E:P combination. | Estimated by fitting to data. |
| $K_{D3}$ | Baseline E:G binding. Different for every E:G combination. | Fixed to either reported or approximated values. |
| $\alpha$ | Cooperativity of MG binding. | Estimated by fitting to data. |
| $\delta$ | Scaling of binding affinities due to different in assay conditions (Yamanaka et al. data only) | Estimated by fitting to data. |
| $P_T$ | Abundance of target protein. Different for each assay. During iMRM experiment simulation, also different for each target protein. | Estimated by fitting to data. |
| $E_T$ | Abundance of effector protein. Different for each assay. | Estimated by fitting to data. |
| $\gamma$ | Baseline degradation rate constant for target proteins. Assumed to be equal for all target proteins as simplifying assumptions | Estimated by fitting to data. |
| $k_{cat}$ | Ternary complex induced degradation rate constant. Depends on all binding partners in EGP complex | Estimated by fitting to data. |

**Supplementary Table S4:** Description of literature sources for data used in model validation.

| Source | Modality | Effector Proteins | Molecular Glues | Target Proteins | Assays Used |
| --- | --- | --- | --- | --- | --- |
| Deutscher et al. | Stabilizer | FKBP12 | Cmpd5,6,7 | FRB | Fluorescence polarization |
| Schulze et al. | Stabilizer | CYPA | Cmpd-3, 4, RMC-4998 | KRAS | TR-FRET, MS |
| Vickery et al. | Stabilizer | 14-3-3 | CRAF01-04, TAZ01-03, ERa01-02 | CRAF, TAZ, ERa | BRET |
| Sperling et al. | Degrader | CRBN | Lenalidomide, Pomalidomide, Avadomide, cc885 | IKZF1, IKZF3, ZNF692, ZFP91, RNF166, GSPT1 | BRET, iMRM |
| Yamanaka et al. | Degrader | CRBN | Lenalidomide, Pomalidomide, Thalidomide | IKZF1, SALL4, PLZF | AlphaScreen, HIBIT. |

**Supplementary Table S5:** Estimated model parameters fitting Deutscher et al. dataset. Reported values are mean (CV) for 6 parameter sets, reported to two significant digits. n/a indicates parameter was not estimated for that drug/target combination.

| Parameter | Mean (CV) | Parameter | Mean (CV) |
| --- | --- | --- | --- |
| $K_{D1}^{Cmpd5}$ | 5.5 nM (fixed) | $K_{D2}^{Cmpd5-FRB}$ | 1000uM (1e-5) |
| $K_{D1}^{Cmpd6}$ | 86 nM (fixed) | $K_{D2}^{Cmpd6-FRB}$ | 620 uM (1e-5) |
| $K_{D1}^{Cmpd7}$ | 6.3 nM (fixed) | $K_{D2}^{Cmpd7-FRB}$ | 77uM (1e-5) |

**Supplementary Table S6:** Estimated model parameters fitting Schulze et al. dataset. Reported values are mean (CV) for 9 parameter sets, reported to two significant digits. n/a indicates parameter was not estimated for that drug/target combination.

| Parameter | Target Protein |  |  |
| --- | --- | --- | --- |
|  | Cmpd-3 | Cmpd-4 | RMC-4998 |
| $k_{b1} (s^{-1})$ | 0.24 (2.0) | 0.42 (1.7) | 0.0011 (0.12) |
| $k_{b2} (s^{-1})$ | 0.61 (2.2) | 0.029 (1.0) | 0.001 (2.1e-6) |
| $k_{cat} (s^{-1})$ | 0.0079 (3.0) | 0.0034 (0.64) | 0.14 (0.00062) |

**Supplementary Table S7:** Estimated model parameters fitting Vickery et al. dataset. Reported values are mean (CV) for 28 parameter sets, reported to two significant digits. n/a indicates parameter was not estimated for that drug/target combination.

| CRAF |  | TAZ |  | ERa |  |
| --- | --- | --- | --- | --- | --- |
| Parameter | Mean (CV) | Parameter | Mean (CV) | Parameter | Mean (CV) |
| $E_T$ | 170 nM (0.031) | $E_T$ | 73 nM (0.0064) | $E_T$ | 72 nM (0.13) |
| $P_T$ | 1 nM (0.011) | $P_T$ | 1 nM (9.0e-5) | $P_T$ | 70 nM (0.13) |
| $K_{D1}^{CRAF}$ | 100 nM (0.030) | $K_{D1}^{TAZ}$ | 100 nM (8.2e-5) | $K_{D1}^{ERa}$ | 110 nM (0.13) |
| $\alpha^{CRAF01-CRAF}$ | 1.0 (0.0088) | $\alpha^{TAZ01-TAZ}$ | 1.1 (0.0018) | $\alpha^{ERa01-ERa}$ | 30 (0.022) |
| $\alpha^{CRAF02-CRAF}$ | 87 (0.084) | $\alpha^{TAZ02-TAZ}$ | 33 (0.056) | $\alpha^{ERa02-ERa}$ | 1.2 (0.0059) |
| $\alpha^{CRAF03-CRAF}$ | 6.3 (0.0089) | $\alpha^{TAZ03-TAZ}$ | 3.3 (0.0071) | $K_{D3}^{ERa01}$ | 10000 nM (0.0038) |
| $\alpha^{CRAF04-CRAF}$ | 3.6 (0.0038) | $K_{D3}^{TAZ01}$ | 100 nM (3.0e-5) | $K_{D3}^{ERa02}$ | 100 nM (0.012) |
| $K_{D3}^{CRAF01}$ | 510 nM (3.9) | $K_{D3}^{TAZ01}$ | 9900 nM (0.057) | | |
| $K_{D3}^{CRAF02}$ | 9700 nM (0.084) | $K_{D3}^{TAZ01}$ | 100 nM (1.1e-5) | | |
| $K_{D3}^{CRAF03}$ | 690 nM (0.011) | | | | |
| $K_{D3}^{CRAF04}$ | 1300 nM (0.0087) | | | | |

**Supplementary Table S8:** Estimated model parameters fitting Sperling et al. dataset. Reported values are mean (CV) for 29 parameter sets, reported to two significant digits. n/a indicates parameter was not estimated for that drug/target combination.

| Parameter | Target Protein |  |  |  |  |  |
| --- | --- | --- | --- | --- | --- | --- |
|  | IKZF1 | IKZF3 | GSPT1 | ZFP91 | RNF166 | ZNF692 |
| $K_{D1}(nM)$ | 370 (0.30) | 200 (0.29) | 330 (0.31) | 300(0.30) | 520 (0.30) | 540 (0.30) |
| $\alpha_{len}$ | 43 (0.017) | 30 (0.10) | 4.3 (0.012) | 9.5 (0.0089) | 5.3 (0.31) | 84 (0.012) |
| $\alpha_{pom}$ | 89 (0.018) | 37 (0.12) | 6.7 (0.019) | 55 (0.015) | 6.2 (0.33) | 110 (0.012) |
| $\alpha_{ava}$ | 410 (0.015) | 160 (0.15) | n/a | 1.2e3 (0.021) | 66 (0.59) | 140 (0.011) |
| $\alpha_{cc885}$ | 290 (0.032) | 26 (0.031) | 1.2e5 (2.5) | n/a | 14 (0.59) | 35 (0.012) |
| $k_{cat}^{len} (hr^{-1})$ | 1.4 (0.99) | 1.0 (0.35) | n/a | 0.71 (0.61) | 6.1 (4.5) | 21 (4.7) |
| $k_{cat}^{pom} (hr^{-1})$ | 2.9 (1.2) | 2.4 (0.37) | n/a | 3.3 (0.86) | 6.1 (4.4) | 420 (0.98) |
| $k_{cat}^{ava} (hr^{-1})$ | 230 (1.6) | 770 (0.44) | n/a | 2.9 (1.0) | 7.5 (4.6) | 960 (0.087) |
| $k_{cat}^{cc885} (hr^{-1})$ | 76 (3.0) | 250 (1.6) | 320 (1.3) | n/a | 37 (5.0) | 1.5 (0.33) |
| $P_T^{MRM} (nM)$ | 21 (1.6) | 5.1 (2.0) | 0.094 (3.0) | 33 (1.3) | 32 (1.1) | 4.5 (2.2) |
| $\gamma (hr^{-1})$ | 0.064 (0.22) | | | | | |
| $P_T^{BRET} (nM)$ | 62 (0.30) | | | | | |
| $E_T^{BRET} (nM)$ | 0.12 (1.8) | | | | | |
| $E_T^{MRM} (nM)$ | 3.4 (0.27) | | | | | |

**Supplementary Table S9:**
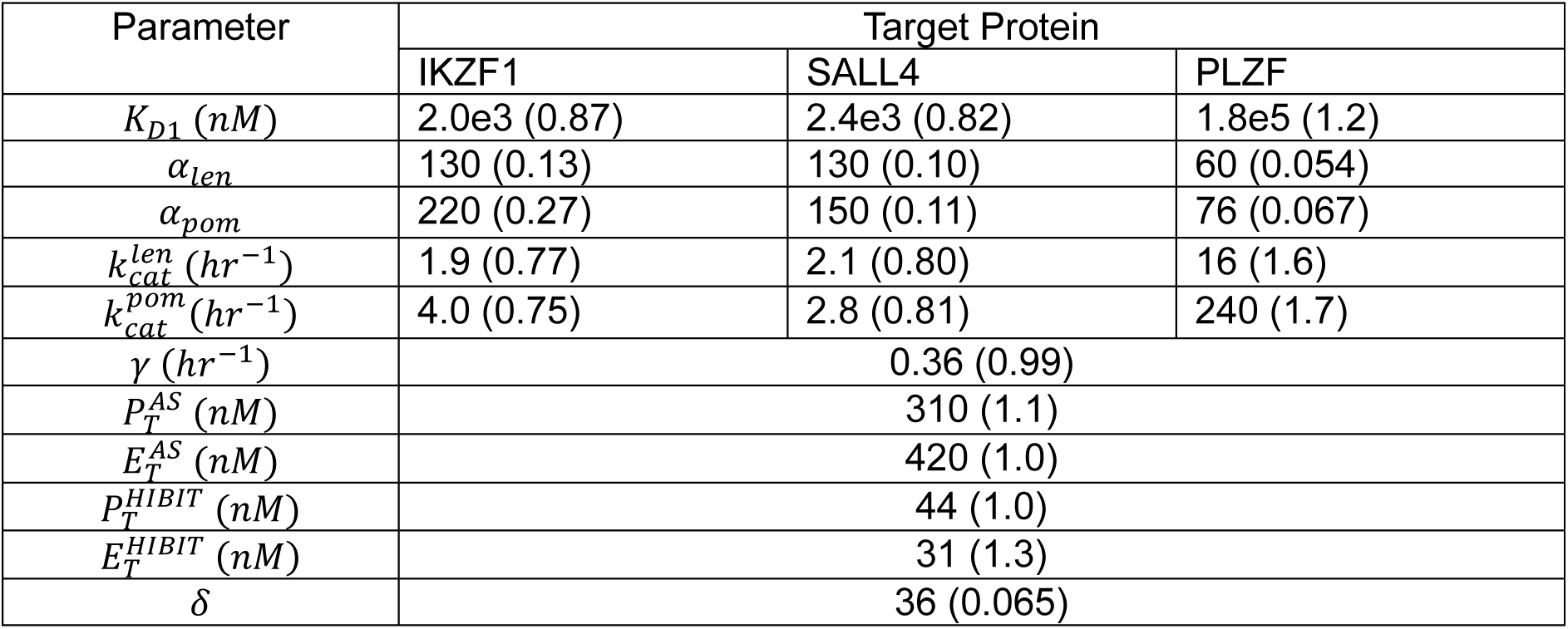
Estimated model parameters fitting Yamanaka et al. dataset. Reported values are mean (CV) for 10 parameter sets, reported to two significant digits.

| Parameter | Target Protein |  |  |
| --- | --- | --- | --- |
|  | IKZF1 | SALL4 | PLZF |
| $K_{D1} (nM)$ | 2.0e3 (0.87) | 2.4e3 (0.82) | 1.8e5 (1.2) |
| $\alpha_{len}$ | 130 (0.13) | 130 (0.10) | 60 (0.054) |
| $\alpha_{pom}$ | 220 (0.27) | 150 (0.11) | 76 (0.067) |
| $k_{cat}^{len} (hr^{-1})$ | 1.9 (0.77) | 2.1 (0.80) | 16 (1.6) |
| $k_{cat}^{pom} (hr^{-1})$ | 4.0 (0.75) | 2.8 (0.81) | 240 (1.7) |
| $\gamma (hr^{-1})$ | 0.36 (0.99) | | |
| $P_T^{AS} (nM)$ | 310 (1.1) | | |
| $E_T^{AS} (nM)$ | 420 (1.0) | | |
| $P_T^{HIBIT} (nM)$ | 44 (1.0) | | |
| $E_T^{HIBIT} (nM)$ | 31 (1.3) | | |
| $\delta$ | 36 (0.065) | | |

**Supplementary Table 10:** Best-fitting model parameter set estimated via fit to Vickery et al. data used for generation of MG stabilizer analysis figures. Values reported to two significant digits.

| CRAF |  |
| --- | --- |
| Parameter | Mean (CV) |
| $E_T$ | 170 nM |
| $P_T$ | 1 nM |
| $K_{D1}^{CRAF}$ | 100 nM |
| $\alpha^{CRAF01-CRAF}$ | 1 |
| $\alpha^{CRAF02-CRAF}$ | 89 |
| $\alpha^{CRAF03-CRAF}$ | 6.3 |
| $\alpha^{CRAF04-CRAF}$ | 3.6 |
| $K_{D3}^{CRAF01}$ | 100 nM |
| $K_{D3}^{CRAF02}$ | 10000 nM |
| $K_{D3}^{CRAF03}$ | 690 nM |
| $K_{D3}^{CRAF04}$ | 1300 nM |

**Supplementary Table 11:**
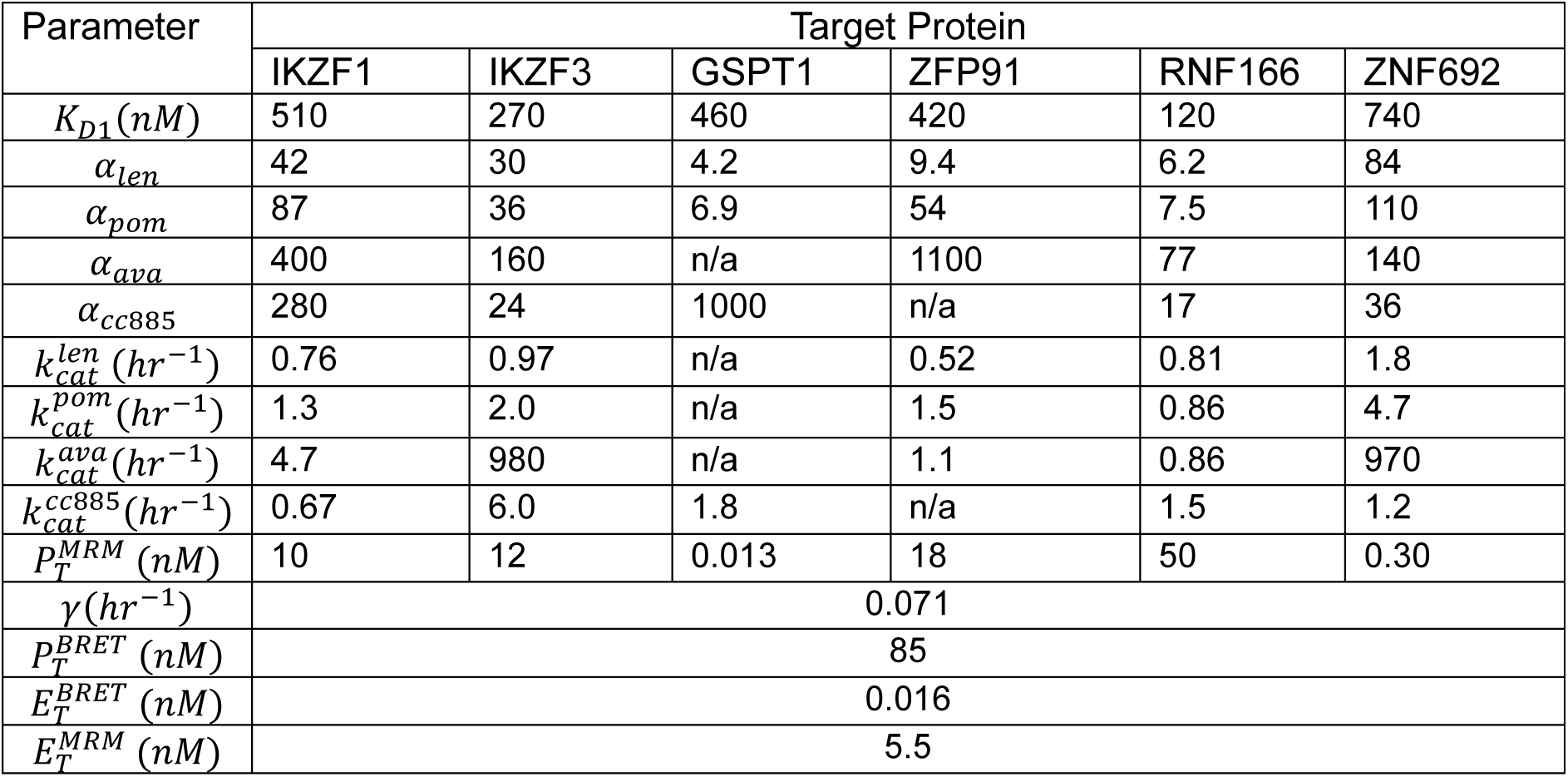
Best-fitting model parameter set estimated via fit to Sperling et al. data used for generation of degrader analysis figures. Values reported to two significant digits.

## Supplementary Figures

**Supplementary Figure 1:**
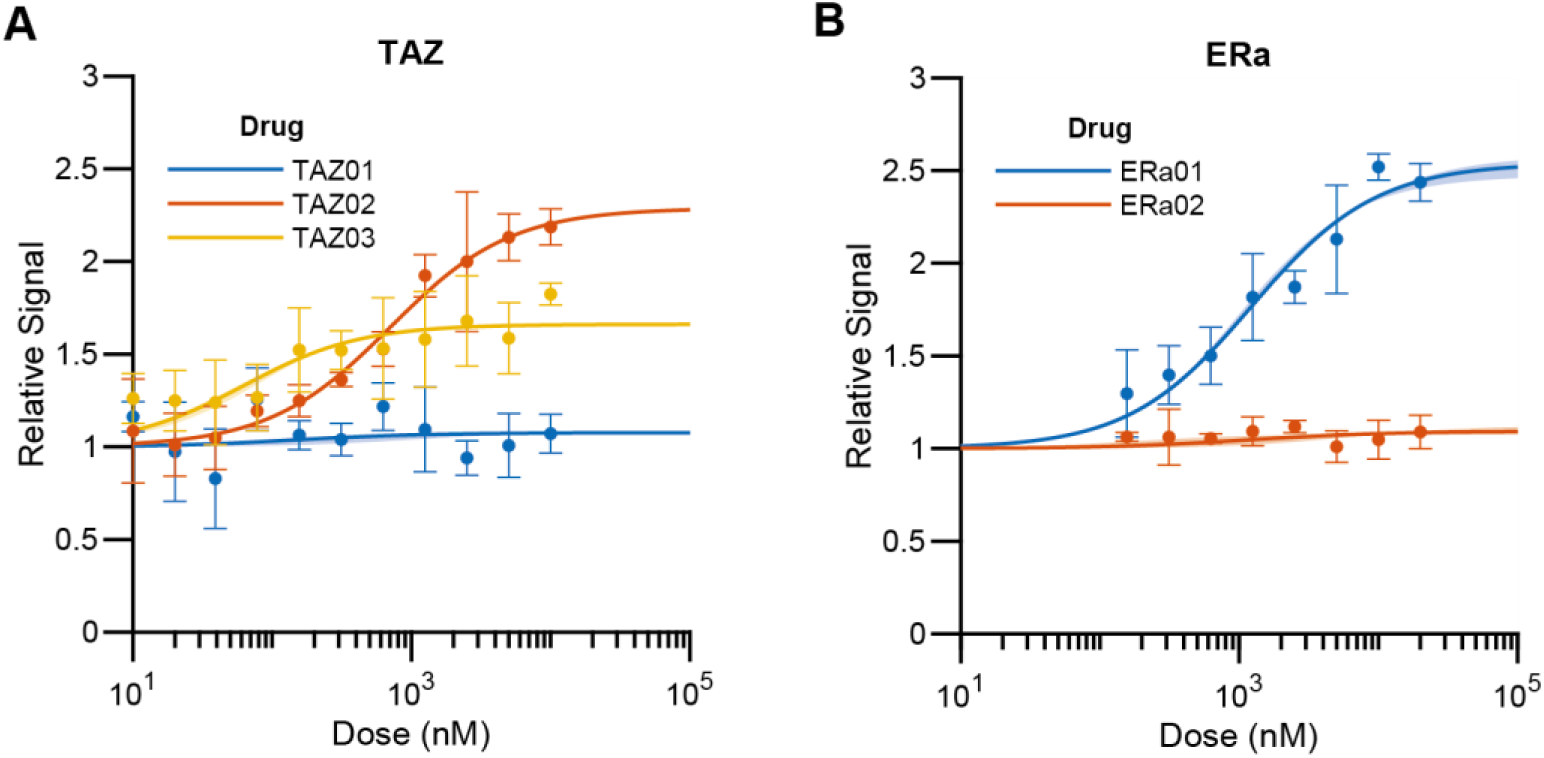
Fitted MG stabilizer model predictions compared to nanoBRET dataset by Vickery et al.^1^. (A) Ensemble (n=16) model fitness, based on the model in Figure 1C, describing ternary complex (14-3-3:MG:TAZ) formation induced by TAZ01 (blue), TAZ02 (red), and TAZ03 (yellow). (B) Ensemble (n=11) model fitness, based on the model in Figure 1C, describing ternary complex (14-3-3:MG:ERa) formation induced by ERa1 (blue) and ERa02 (red). Solid curves: Mean predicted values. Shaded regions: Raw 95% interval. Points: experimental data.

**Supplementary Figure 2:**
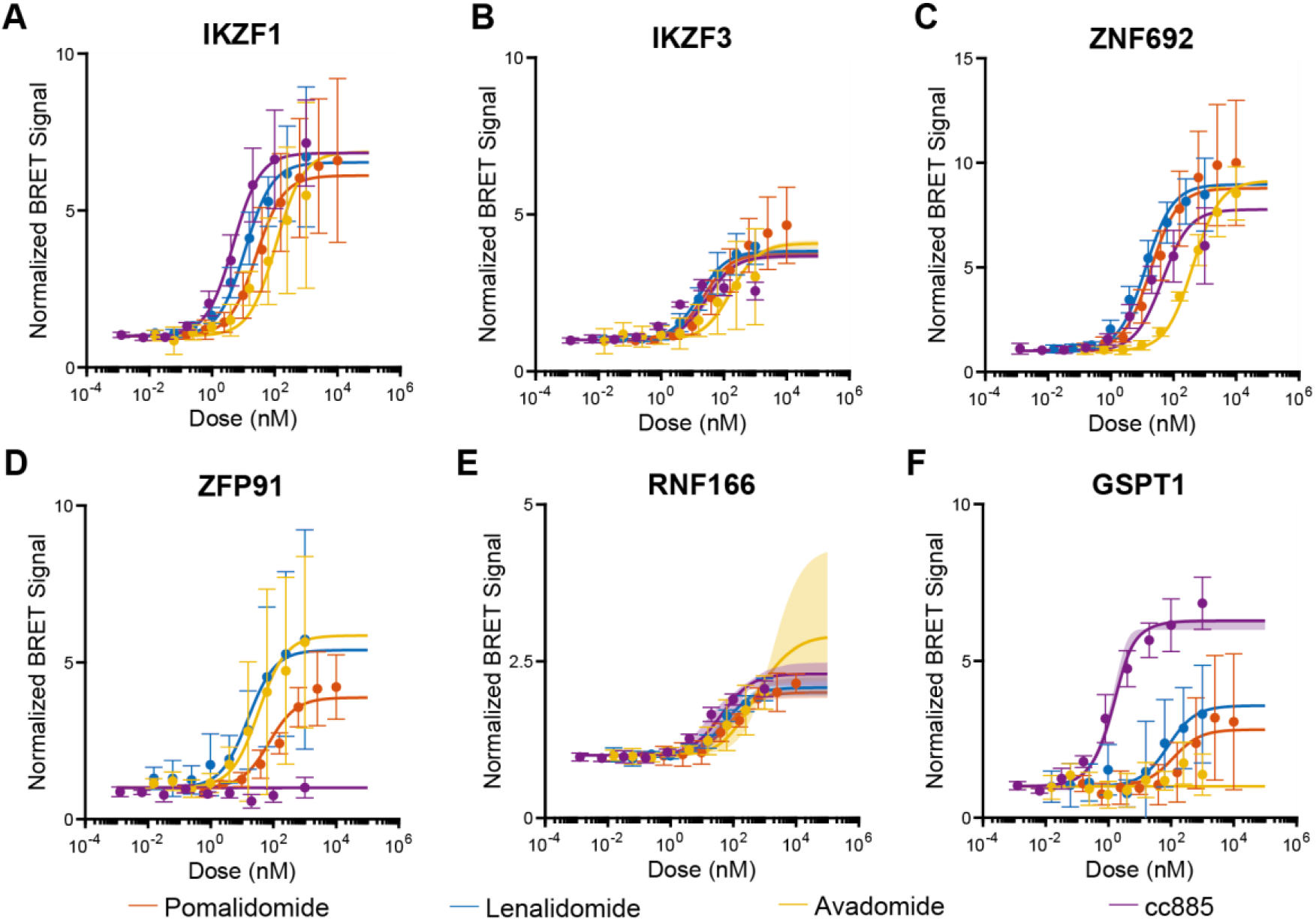
Fitted MG degrader model predictions compared to BRET dataset presented by Sperling et al.^2^. Ensemble (n = 29) model fitness to BRET data for A) IKZF1, B) IKZF3, C) ZNF692, D) ZFP91, E) RNF166, F) GSPT1. Solid curves: mean predicted value. Shaded region: Raw 95% interval. Orange: Pomalidomide. Blue: Lenalidomide. Yellow: Avadomide. Purple: cc885.

**Supplementary Figure 3:**
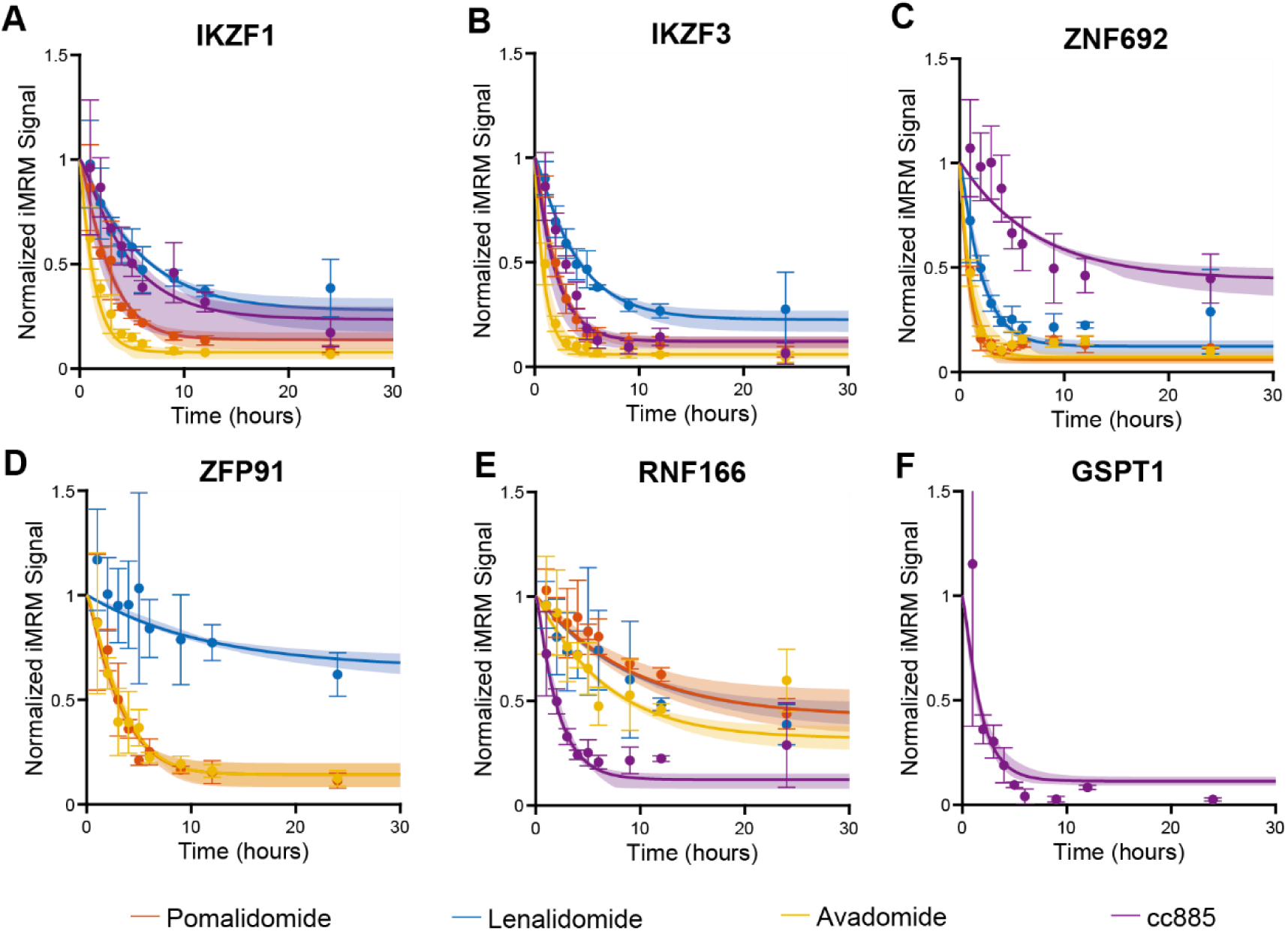
Fitted MG degrader model predictions compared to iMRM timecourse dataset presented by Sperling et al.^2^. Ensemble (n = 29) model fitness to iMRM timecourse data for A) IKZF1, B) IKZF3, C) ZNF692, D) ZFP91, E) RNF166, F) GSPT1. Solid curves: mean predicted value. Shaded region: Raw 95% interval. Orange: Pomalidomide. Blue: Lenalidomide. Yellow: Avadomide. Purple: cc885.

**Supplementary Figure 4:**
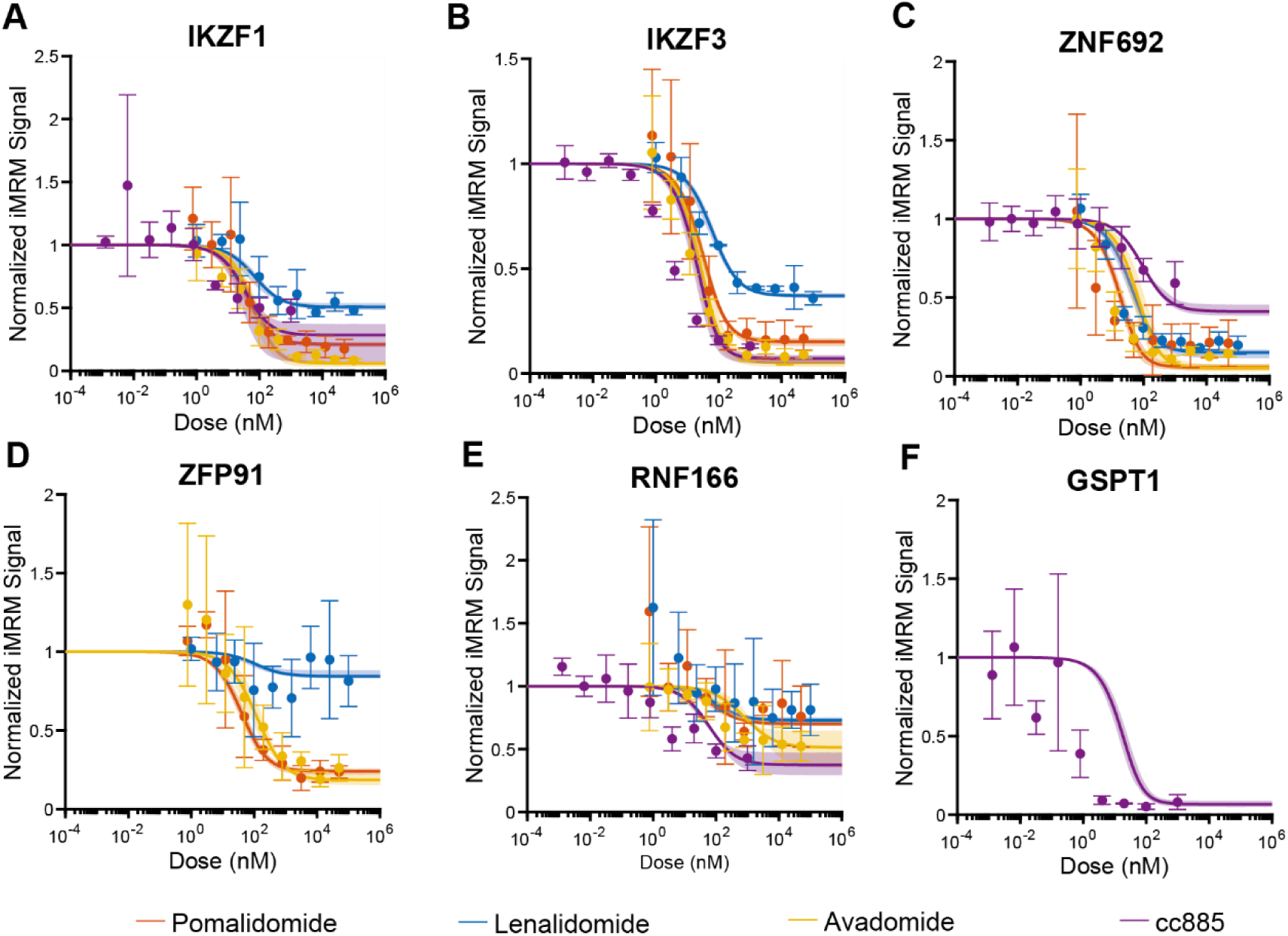
Fitted MG degrader model predictions compared to iMRM dose response dataset presented by Sperling et al.^2^. Ensemble (n = 29) model fitness to iMRM dose response data for A) IKZF1, B) IKZF3, C) ZNF692, D) ZFP91, E) RNF166, F) GSPT1. Solid curves: mean predicted value. Shaded region: Raw 95% interval. Orange: Pomalidomide. Blue: Lenalidomide. Yellow: Avadomide. Purple: cc885.

**Supplementary Figure 5:**
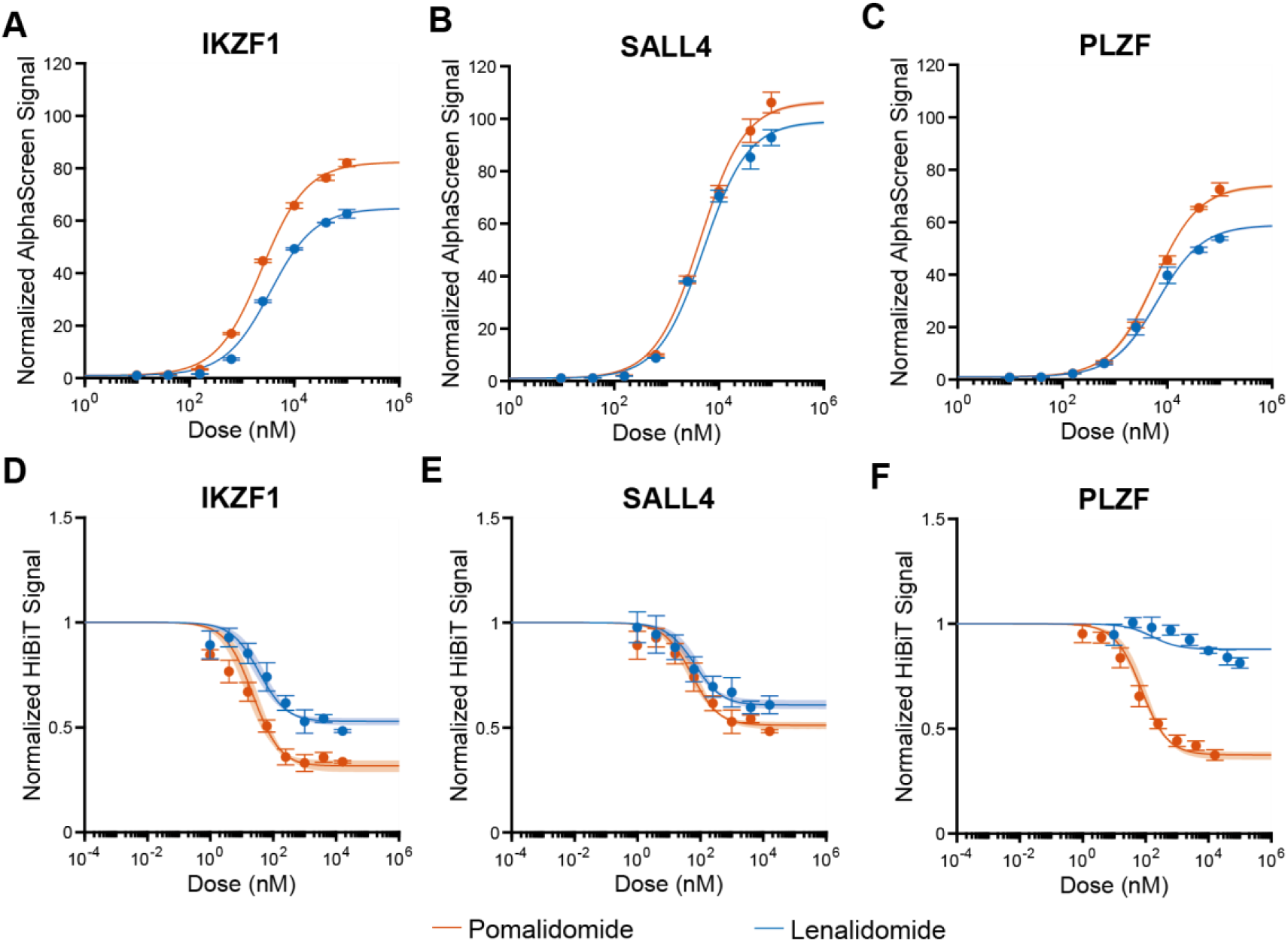
Fitted MG degrader model predictions compared to datasets presented by Yamanaka et al.^3^. A-C) Ensemble (n = 10) model predictions compared to A) IKZF1, B) SALL4, and C) PLZF AlphaScreen titrations. D-F) Ensemble (n = 10) model predictions compared to A) IKZF1, B) SALL4, and C) PLZF HiBiT titrations. Solid curves: Mean predicted value. Shaded region: Raw 95% interval. Orange: Pomalidomide. Blue: Lenalidomide.

